# Restoration of Capacity to Build Muscle Strength in Geriatric Mice by Inhibition of the Gerozyme 15-Prostaglandin Dehydrogenase

**DOI:** 10.64898/2026.08.06.743387

**Authors:** Minas Nalbandian, Ireh Kim, Elena Monti, Yutong Kelly Li, Emmeran Le Moal, Peggy E. Kraft, Kaitlin Jeuris, Michelle To, Ludmila Alexandrova, Jameel Barkat, Zeyuan Zhang, Katrin Svensson, Helen M. Blau

## Abstract

Loss of skeletal muscle mass and strength with age drives sarcopenia, a syndrome affecting >100 million people worldwide that leads to loss of mobility, independence, and increased mortality. Mechanical overload induces hypertrophy in young muscle, but this response is markedly attenuated with age—a poorly understood phenomenon termed “anabolic resistance.” Here we test whether impaired paracrine communication between myofibers and their niche underlies this loss of plasticity in geriatric mice. In aged muscle, pharmacological inhibition of 15-PGDH restores prostaglandin E2 (PGE2) bioavailability and rescues the anabolic response, increasing muscle growth and contractile strength. Single-nuclei RNA-seq revealed a paracrine circuit: PGE2 drives IGF1 synthesis in type IIb myonuclei, which signals to stromal, myogenic, myonuclear, and immune cells. Blocking IGF1 receptor signaling abolished these gains, placing PGE2 upstream of an IGF1-mediated circuit that coordinates multicellular hypertrophy. Thus, 15-PGDH inhibition is a pharmacological strategy to overcome the anabolic resistance and rebuild muscle in aging.

## Introduction

Skeletal muscle mass and strength are critical determinants of mobility, metabolic health, and quality of life. Both decline steadily with age. Humans lose ∼10% of their skeletal muscle mass per decade after age 50^1^. This progressive loss can culminate in sarcopenia, a disabling clinical syndrome affecting approximately 10–27% of adults aged 60 years and older^2–5^. Despite its prevalence, sarcopenia has only recently been recognized by an ICD code (M62.84) and a consensus of diagnostic criteria based on validated clinical measures^3,4,6^. Clinically, severe muscle wasting and weakness increase the risk of falls and fractures, accelerate loss of independence, and are associated with higher mortality^7,8^, creating a growing public health and economic burden worldwide^9^.

Although sarcopenia is multifactorial — arising from a complex combination of endocrine changes, chronic inflammation, loss of neuromuscular connectivity, and reduced physical activity—a key limiting feature is the reduced capacity of aged muscle to adapt to anabolic stimuli. This phenomenon, often termed anabolic resistance^10^, reflects an impaired translation of stimuli such as mechanical loading, nutrient and hormonal signals into increases in muscle protein and improvement in function. Consistent with this definition, after ∼6 weeks of resistance training, young individuals exhibit ∼20% increases in strength, whereas older cohorts performing the same exercise regimen typically achieve only ∼5% gains^11^. This difference underscores the markedly blunted hypertrophic response to mechanical loading that occurs with aging. The gap between clinical need and biological responsiveness is particularly acute during recovery from illness, injury, or disuse, when muscle atrophy is particularly severe. To date there are no approved therapies that meet this need.

Mechanical overload, as induced by resistance training, initiates a tightly coordinated cascade of molecular and cellular events, including calcium influx, sarcolemmal deformation, and activation of kinase signaling pathways^12,13^, which converge on anabolic networks such as mTORC1^14–16^. Together, these programs activate muscle stem cells, drive myofiber hypertrophy, and remodel the extracellular matrix, thereby increasing both muscle mass and strength^17–20^. In young muscle, this machinery is highly responsive; in aged muscle, however, even strong overload stimuli result in limited growth and force gains^21,22^. This decline in responsiveness highlights a pressing need to define the molecular basis of anabolic resistance with aging and identify strategies that can overcome it.

Although intrinsic defects in myofibers and muscle stem cells are known to accompany aging, we reasoned that cell-cell interactions and paracrine signaling among non-muscle cell types could be a critical mediator of hypertrophy. Whereas secreted protein growth factors have been extensively characterized and implicated in muscle growth, the role of lipid metabolites has only recently been appreciated. The metabolite Prostaglandin E2 (PGE2) was shown recently to act directly on muscle stem cells^23^, myofibers^24^ and the motoneurons^25^ that innervate them. However, the possibility that PGE2 could act on non-muscle cells within the tissue and impact muscle function indirectly has not yet been explored.

Here, we investigate in geriatric mice (28-month-old)^26,27^ if anabolic resistance can be overcome by pharmacological inhibition of 15-PGDH, the gerozyme that increases with aging and limits the bioavailability of PGE2^24^, and whether this occurs via a paracrine mechanism. We used a well-established mechanical overload model^28^. We find that aging is marked by impaired PGE2 signaling during overload, and that increasing PGE2 bioavailability by 15-PGDH inhibition (PGDHi) rescues muscle mass and contractile force. Type IIb myofibers, which are the first to lose function in aging and are crucial to strength and mobility^29^, become a prominent source of IGF1 in response to PGDHi. snRNA-seq provides support for a paracrine model in which PGE2 induced IGF1 coordinates growth responses across stromal, myogenic, myonuclear, and immune compartments. Strikingly, blockade of IGF1 receptor (IGF1R) signaling in this context prevents the gains in muscle mass and strength, demonstrating that IGF1 is required downstream of PGE2 for the full anabolic response. Together, these findings identify a PGE2–IGF1 paracrine axis as a druggable target for restoring muscle strength and the hypertrophic response in aging skeletal muscle.

## Results

### Muscle hypertrophic capacity declines with aging and is associated with impaired PGE2 signaling

Muscle growth is a fundamental and conserved adaptive process in mammals, enabling structural and functional changes in response to increased physiological demand. However, this capacity declines with aging. To model anabolic resistance in vivo, we used a well-characterized mechanical overload paradigm^28,30–32^ involving tenotomy of the Achilles tendon, which leaves the gastrocnemius and soleus muscles anatomically intact but functionally uncouples them from the ankle, preventing plantarflexion force transmission and redistributing load to the adjacent plantaris muscle. As a result, the plantaris undergoes robust hypertrophy without direct injury, ensuring that the observed adaptations reflect hypertrophic remodeling rather than a regenerative response.

To validate the suitability of this model for studying age-related impairments in muscle hypertrophy, we compared the hypertrophic response of 2-month-old (young) and 28-month-old (geriatric) C57BL/6 male mice. Plantaris muscles were harvested at 2, 4, 6, 7, 9, and 14 days post-tenotomy (**Fig. 1a**). Both young and geriatric mice exhibited progressive increases in plantaris mass following overload, with growth plateauing by day 9 (**Fig. 1b**). However, young animals achieved significantly greater absolute and relative (sham-normalized) plantaris weights than their geriatric counterparts (**Fig. 1b, c**), confirming an age-dependent decline in hypertrophic capacity.

**Figure 1.**
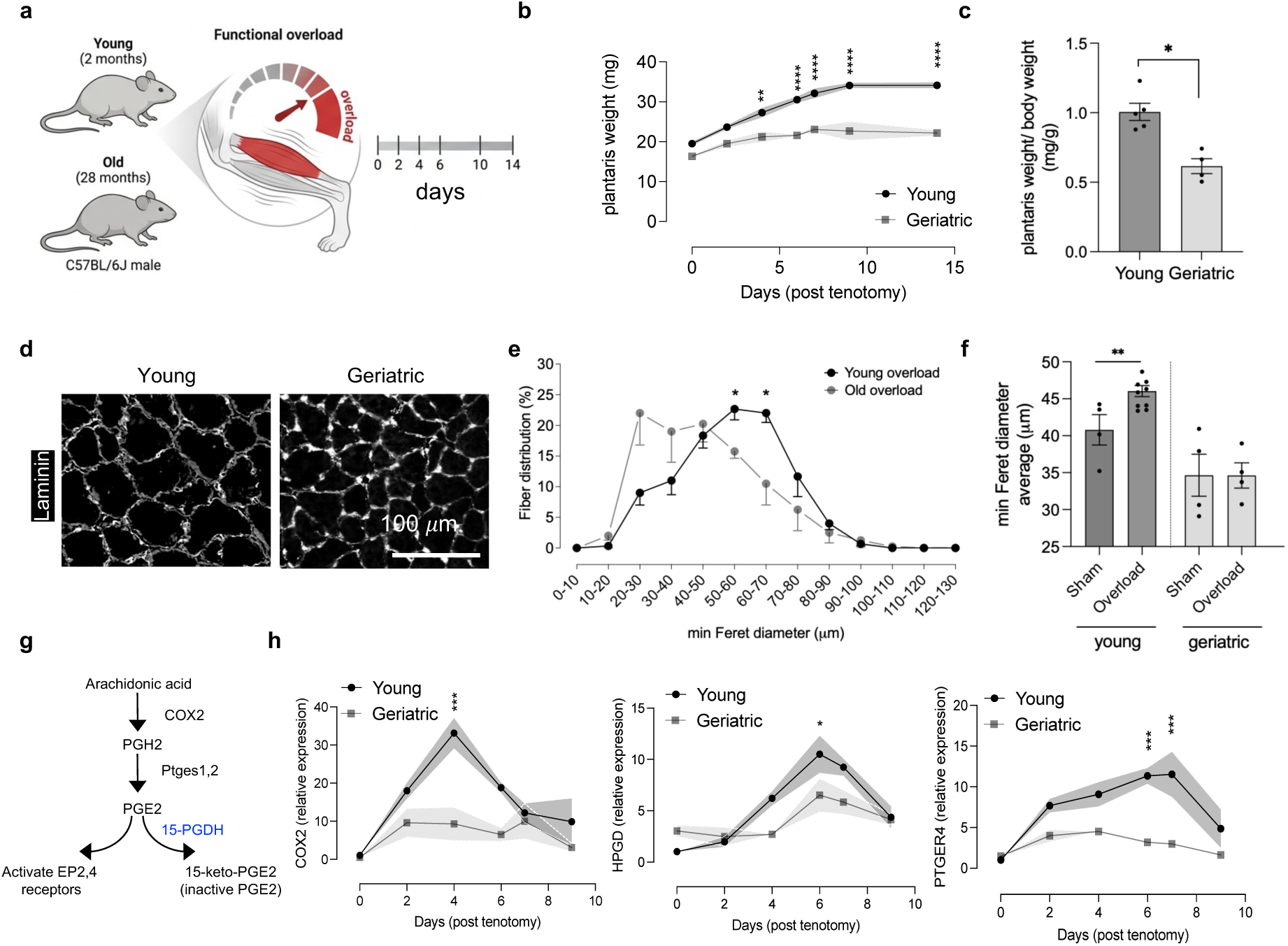
Age-associated decline in overload-induced muscle hypertrophy and PGE2 signaling. (a) Schematic of experimental design in young (2-month-old) and geriatric (28-month-old) C57BL/6 male mice subjected to synergist ablation–induced overload. (b) Time course of plantaris muscle weight following overload (n = 4 per group). (c) Plantaris weight normalized to contralateral sham (n = 4 per group). (d) Representative laminin-stained plantaris cross-sections from young and geriatric mice. Scale bar, 100 μm. (e) Fiber size distribution curves of plantaris myofibers following overload. (f) Quantification of minimal Feret diameters in sham and overloaded plantaris muscles. (g) Schematic of the prostaglandin E2 biosynthetic and degradative pathway. (h) Expression of *Cox2*, *Hpgd* (15-PGDH), and *Ptger4* measured by RT–qPCR at indicated days post-overload (n = 4 per group). In (c) and (f), each dot represents one animal. In (b), (e), and (h), statistical significance indicates comparisons between geriatric and young animals at each time point. Data represent mean ± SEM. \**P* < 0.05, \*\**P* < 0.01, \*\*\**P* < 0.001, **** *P* < 0.0001 by two-way ANOVA with post hoc test or unpaired *t*-test.

As another measure of muscle mass, we assessed myofiber Feret diameter by histological analysis of laminin-stained sections. The average myofiber diameters were significantly larger in overloaded young mice compared to geriatric mice (**Fig. 1d**). Quantification of minimal Feret diameters confirmed that geriatric muscles failed to exhibit hypertrophic growth under overload (**Fig. 1e, f**), demonstrating a blunted adaptive response. These findings indicate that the differences observed under overload reflects in part impaired growth capacity in geriatric muscles.

Muscle stem cells (MuSC) are known to contribute to muscle growth in response to a mechanical stimulus^30,32,33^. To explore MuSC activity as potential cellular drivers of this deficit, we assessed the expression of myogenic genes with a role in regeneration by RT–qPCR. Remarkably, markers of muscle stem cell activation (*Pax7*, *Myf5*, *Myog*) and myoblast fusion (*Mymx*, *Mymk*), as well as *Igf1*, a key anabolic growth factor that promotes muscle hypertrophy and regeneration, were upregulated in young but not geriatric mice (**Fig. S1a**), suggesting that age-related hypertrophic deficits may reflect both impaired muscle stem cell recruitment and defective myonuclear accretion.

Given the established role of prostaglandin E2 (PGE2) in muscle regeneration^23,34^, we interrogated its biosynthetic and degradative pathways in response to overload (**Fig. 1g**). RT–qPCR revealed that overload robustly induced cyclooxygenase-2 (*Cox2*), the rate-limiting enzyme for PGE2 synthesis, with a several-fold higher increase in young compared to geriatric mouse muscle (**Fig. 1h**). Expression of 15-hydroxyprostaglandin dehydrogenase (*15-Pgdh*), which catalyzes PGE2 inactivation, also increased post-overload but peaked one day after *Cox2* induction, suggesting that in young muscles the degrading enzyme is induced as part of a feedback loop to curtail PGE2 signaling. Notably, expression of *Ptger4*, encoding the PGE2 receptor EP4—the predominant receptor isoform in skeletal muscle^24,34,35^—was induced exclusively in young mice (**Fig. 1h**), whereas EP1, EP2, and EP3 receptor transcripts were undetectable (Ct > 32). These findings indicate a dual defect in PGE2 signaling in muscles of geriatric mice: reduced PGE2 production and impaired receptor availability.

### Pharmacological Inhibition of PGE2 Biosynthesis Attenuates Hypertrophy in Young but Not Geriatric Muscle

We wished to determine the basis for the difference in anabolic response to overload in young and aged mice. We hypothesized that de novo PGE2 synthesis might be required for the increase in muscle mass, as seen in young muscles in response to mechanical overload. To test this possibility, young (2-month) and geriatric (28-month) mice were subjected to overload and treated daily with indomethacin, a well-characterized nonsteroidal anti-inflammatory drug (NSAID) that blocks COX-2, the first enzyme required for PGE2 biosynthesis^34^ (**Fig. 2a**).

**Figure 2.**
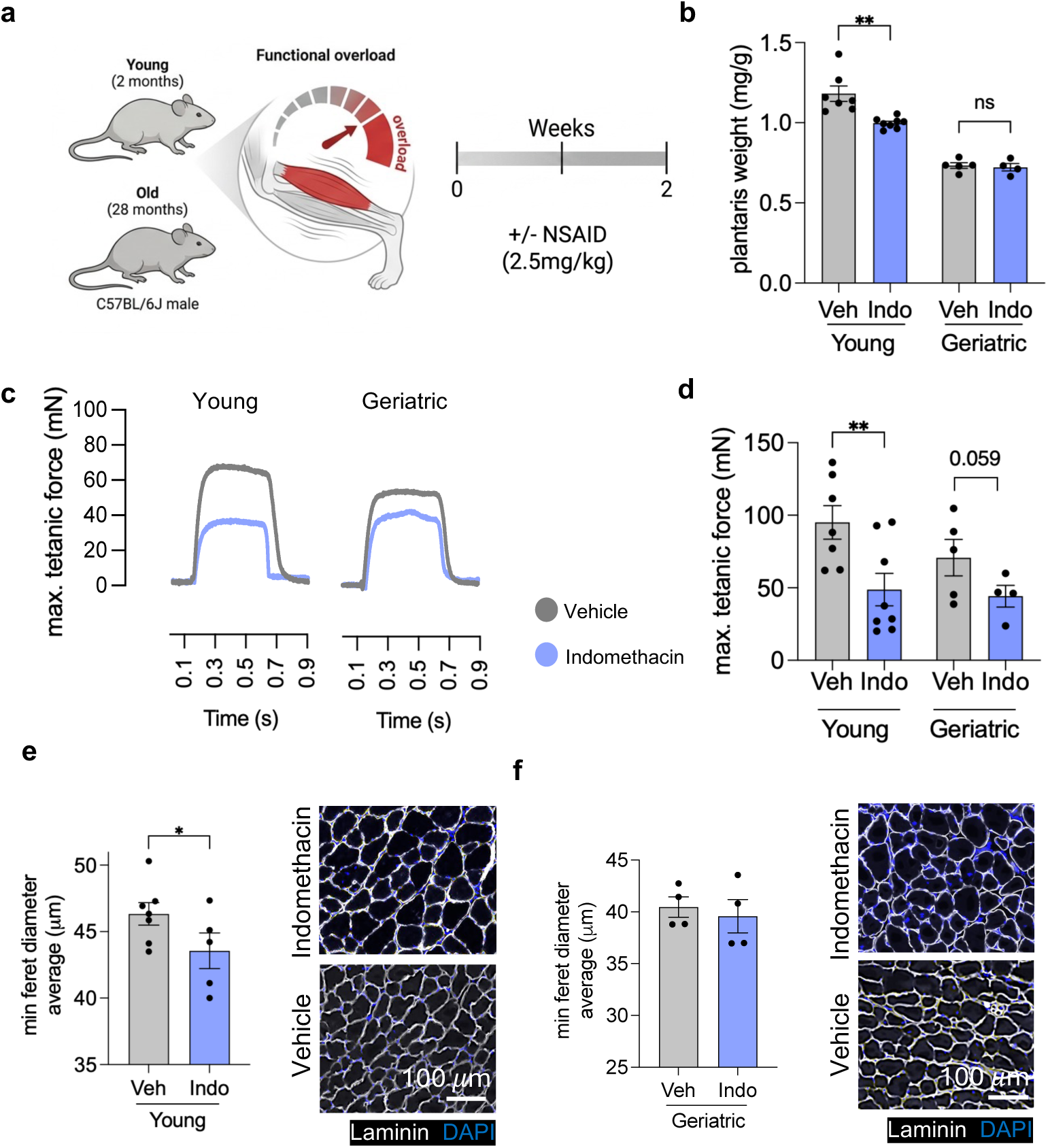
Effect of COX inhibition on overload-induced hypertrophy in young and geriatric mice. (a) Experimental schematic: young (2-month-old) and geriatric (28-month-old) mice subjected to overload with daily intraperitoneal injection of vehicle or indomethacin (2.5 mg/kg). (b) Plantaris muscle weight after 2 weeks of overload (n = 4-7 per group). (c) Representative traces of tetanic force measurements in young and geriatric mice treated with vehicle or indomethacin. (d) Quantification of maximal tetanic force (n = 4-7 per group). (e) Minimal Feret diameter and representative laminin-stained cross-sections of plantaris muscle from young mice treated with vehicle or indomethacin. Scale bar, 100 μm. (f) Minimal Feret diameter and representative laminin-stained cross-sections of plantaris muscle from geriatric mice treated with vehicle or indomethacin. Scale bar, 100 μm. In (b), (d), (e) and (f) each dot represents one animal. Data represent mean ± SEM. Exact *P* values are shown, determined by unpaired *t*-test.

To assess strength, we performed plantar flexion torque analyses. This assay is advantageous because the muscle remains intact, and muscle force is determined by an objective measurement of foot pedal displacement in response to neural stimulation in intact, anesthetized mice —making it less confounded by motivation, learning, or limb use than voluntary tests such as grip strength^24,25,36,37^. After two weeks of overload, indomethacin significantly reduced plantaris weight in young mice compared to vehicle-treated controls, whereas no difference was observed in geriatric animals (**Fig. 2b**). Indomethacin also blunted the overload-induced increase in tetanic force to a greater extent in young mice than in geriatric mice (**Fig. 2c, d**). These data suggest that PGE2 synthesis plays a crucial role in the force-generating response to overload in both young and geriatric mice. Notably, indomethacin’s failure to affect muscle mass while still impairing force in geriatric mice suggests that PGE2 signaling mediates more than hypertrophy alone; other mechanisms, such as an effect on neuromuscular junctions in overloaded geriatric muscle, cannot be ruled out.

Sham-operated muscles showed no changes in plantaris weight or tetanic force with indomethacin treatment (**Fig. S2a, b**), indicating that the detrimental effects of COX-2 inhibition are specific to the overloaded condition. Together, these data indicate that PGE2 plays a key role in the hypertrophic response in both young and aged muscle.

Histological analyses further confirmed these trends: indomethacin treatment reduced average myofiber diameter in young mice, while no significant effect was detected in geriatric muscle (**Fig. 2f, g**). Together, these findings demonstrate that COX-2 activity plays a critical role in mediating overload-induced hypertrophy and strength gains, and suggest that the reduced PGE2 bioavailability associated with aging may contribute to the impaired hypertrophy in aged muscles.

### 15-PGDH Inhibition Reverses Age-Related Impairments in Muscle Hypertrophy

We hypothesized that increasing the bioavailability of PGE2 in geriatric mice could overcome anabolic resistance and restore the hypertrophic response to overload. To test this, young and geriatric mice were treated for two weeks post tendon ablation with PGDHi — a small molecule which inhibits 15-PGDH, the PGE2 degrading enzyme (**Fig. 3a**). We used a highly sensitive liquid chromatography mass spectrometry assay (LC-MS/MS) to distinguish the closely related metabolites, PGE2 and PGD2 (**Fig. S3**) and confirmed that 15-PGDH inhibition elevates muscle PGE2 levels ∼3-fold in old overloaded muscles **(Fig. 3b, c)**. Plantar flexion force measurements were performed at the endpoint, followed by plantaris muscle harvest, weighing, and histological analyses. Importantly, PGDHi treatment did not affect overall body weight or heart weight, indicating that the observed effects were specific to skeletal muscle **(Fig. S4a, b)**.

**Figure 3.**
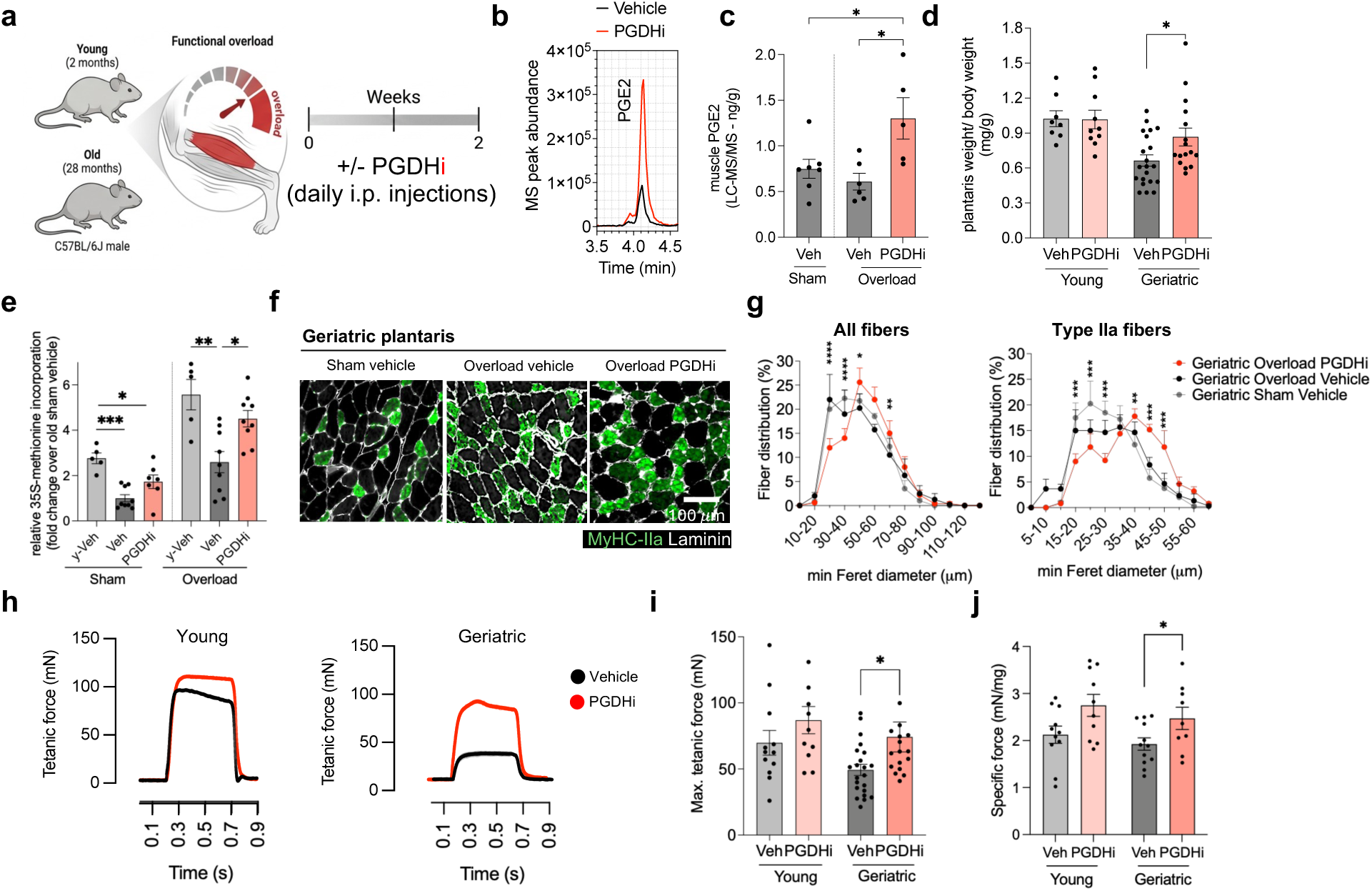
15-PGDH inhibition enhances hypertrophy and function in overloaded geriatric muscle. (a) Experimental schematic of overload-induced hypertrophy in young and geriatric mice treated with vehicle or 15-PGDH inhibitor (PGDHi) for 2 weeks. (b) Representative LC–MS/MS chromatograms of PGE2 levels in overloaded geriatric plantaris muscles treated with vehicle or PGDHi. (c) Muscle PGE2 levels measured by LC–MS/MS in sham and overloaded plantaris muscles (n = 5-7 per group). (d) Plantaris weight normalized to body weight in young and geriatric mice after overload with vehicle or PGDHi treatment (n = 8-21 per group). (e) Levels of protein synthesis measured by [^35^S]-methionine incorporation in plantaris muscles overloaded for four days. (f) Representative laminin (white) and type IIa fiber (MyHC-IIa, green) immunostaining of geriatric plantaris muscle sections from sham and overloaded mice treated with vehicle or PGDHi. Scale bar, 100 μm. (g) (left) Fiber size distribution of all fibers in geriatric overloaded muscles treated with vehicle or PGDHi (n = 4-8 per group). (right) Fiber size distribution of type IIa fibers in geriatric overloaded muscles treated with vehicle or PGDHi (n = 4-8 per group). (h) Representative force traces of plantar flexor tetanic contractions from young and geriatric mice treated with vehicle or PGDHi. (i) Maximal tetanic force of plantaris muscles in young and geriatric mice with vehicle or PGDHi treatment (n = 8-21 per group). (j) Specific force (normalized to muscle weight) in young and geriatric mice treated with vehicle or PGDHi (n = 8-21 per group). In (c), (d), (e), (i) and (j) each dot represents one animal. Data represent mean ± SEM. \**P* < 0.05, \*\**P* < 0.01, \*\*\**P* < 0.001, **** *P* < 0.0001 by unpaired *t*-test.

This increase in PGE2 was accompanied by increased muscle mass after two weeks of PGDHi treatment and elevated protein synthesis, measured by [^35^S]-methionine incorporation, four days after overload induction in geriatric mice (**Fig. 3d, e**). This day 4 time point coincided with peak COX-2 expression and the highest rate of overload-induced muscle growth observed in Figure 1. Consistent with this early anabolic response, PGDHi in geriatric muscles also increased myofiber cross-sectional area relative to vehicle-treated controls two weeks after overload induction (**Fig. 3f, g**). Because type IIa fibers have been reported to be preferentially targeted during early MuSC-mediated fusion events during overload-induced hypertrophy^38^ and represent a growth-responsive fast oxidative fiber population^39^, we next asked whether PGDHi preferentially affected this fiber type. Fiber-type analysis revealed that PGDHi preferentially increased the cross-sectional area of type IIa fibers (**Fig. 3g**).

Functionally, PGDHi treatment significantly enhanced both absolute force production and specific force, the latter reflecting force normalized to muscle mass and serving as a measure of muscle quality in geriatric mice. In contrast, young mice showed only a modest, non-significant trend toward improvement (**Fig. 3h–j**). The striking increase in force in aged overloaded muscles was approximately 50% relative to vehicle controls, nearly five-fold greater than the effect observed with PGDHi alone in sham-operated limbs (**Fig. S4c**). These findings suggest that PGDHi acts synergistically with mechanical overload rather than merely producing an additive effect. Thus, restoration of PGE2 signaling may fundamentally alter how aged muscle responds to anabolic stimulation, positioning 15-PGDH as a limiting factor in age-related anabolic resistance and impaired hypertrophic adaptation.

### 15-PGDH Inhibition Enhances Muscle Stem Cell Activation and Myonuclear Accretion in Geriatric Muscles

We reasoned that the increase in muscle mass and strength during overload induced hypertrophy could result in part from an increase in stem cell activity, even in the absence of injury. We previously showed that PGE2 is required for muscle stem cell proliferation in response to notexin-induced injury^23,34^. To determine whether muscle stem cells contributed to the increase in muscle size during non-injury hypertrophy, we treated young and geriatric mice with the 15-PGDH inhibitor (PGDHi) and administered 5-ethynyl-2’-deoxyuridine (EdU) in drinking water at a concentration of 0.2 mg/mL to label proliferating cells during the early phase of overload-induced growth (**Fig. 4a**). EdU was provided continuously for up to 7 days because this window captures the early proliferative response after mechanical overload, before the later two-week hypertrophic endpoint. Plantaris muscles were harvested at days 2, 5, and 7 post-tenotomy to define the kinetics of cell proliferation and early growth. By day 5 post-tenotomy, geriatric mice treated with PGDHi already showed a marked hypertrophic response that was not observed in vehicle-treated controls (**Fig. 4b**).

**Figure 4.**
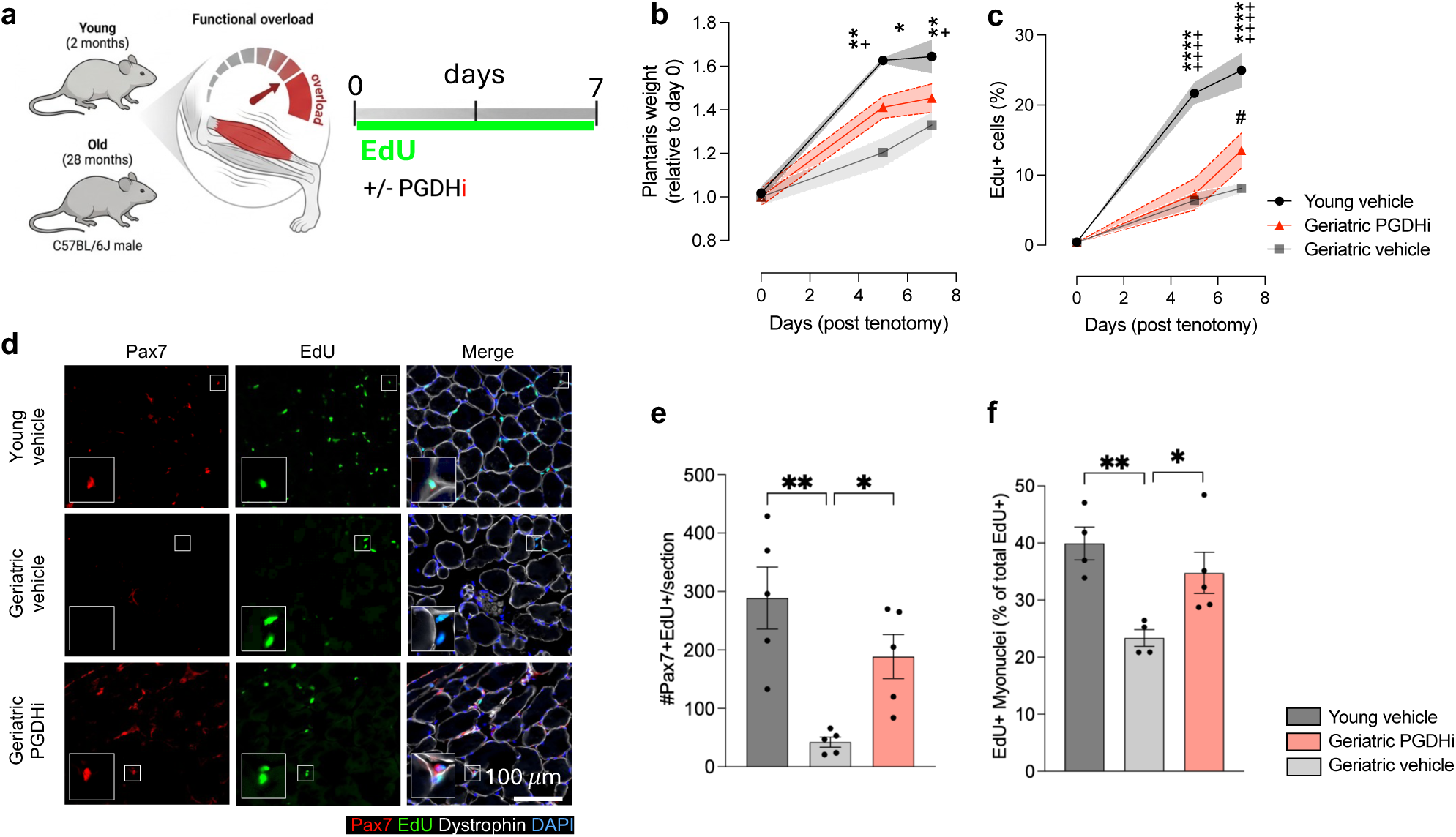
PGDHi enhances muscle stem cell activity and myonuclear accretion in overloaded geriatric muscles. (a) Experimental design: young (2-month-old) and geriatric (28-month-old) mice underwent overload surgery and were treated with vehicle or PGDHi for 7 days, with EdU administered continuously in drinking water. (b) Plantaris muscle weight relative to baseline (n = 4-5 per group). (c) Percentage of EdU⁺ cells over the time course (n = 4-5 per group). (d) Representative immunostaining of Pax7 (red), EdU (green), dystrophin (white), and DAPI (blue) in plantaris muscle sections. Insets highlight Pax7⁺EdU⁺ cells. (e) Number of Pax7⁺EdU⁺ nuclei per section on day 7 after tenotomy (n = 4-5 per group). (f) Number of EdU⁺ myonuclei per section on day 7 after tenotomy (n = 4-5 per group). In (e) and (f) each dot represents one animal. Data represent mean ± SEM. \**P* < 0.05, \*\**P* < 0.01, \*\*\**P* < 0.001, **** *P* < 0.0001 by one-way ore two-way ANOVA with post hoc test.

To evaluate the contribution of stem cells to the hypertrophic response, we performed histological analysis. We found that the total number of EdU+ cells was significantly lower in geriatric mice compared to young controls as expected (**Fig. 4c**). Notably, PGDHi led to a marked increase in EdU+ cell numbers in aged mouse plantaris muscles by day 7. Co-staining for Pax7 and EdU demonstrated fewer proliferating muscle stem cells (Pax7+EdU+) in aged mice relative to young mice, with PGDHi treatment partially rescuing this deficit (**Fig. 4d, e**). We hypothesized that this increase in muscle stem cell proliferation would be accompanied by enhanced fusion of newly generated nuclei into growing myofibers. To test this, we quantified EdU+ nuclei within myofibers as a proxy for newly incorporated myonuclei. Young muscles exhibited more EdU+ myonuclei than geriatric muscles, and PGDHi treatment increased this population in geriatric mice (**Fig. 4f**). Together, these results indicate that PGDHi enhances muscle stem cell proliferation and myonuclear accretion during overload-induced hypertrophy in geriatric muscle, suggesting that stem cells are a major contributor to the increase in muscle function and that PGDHi can partially restore the growth-associated activity of aged muscle stem cells.

### Single-Nuclei Transcriptomic Profiling Reveals PGDHi-mediated Cell Type–Specific Overload Responses in Young and Geriatric Muscles

We sought to investigate the cellular and molecular basis for the differential overload responses in young versus geriatric mouse muscles, and how these responses are modulated by 15-PGDH inhibition (PGDHi). We performed single-nuclei RNA sequencing (snRNA-seq) on plantaris muscles, as muscle is a syncytium and single cell analyses would omit a large proportion of nuclei. We compared snRNA-seq from sham-operated and overloaded (14 days post-tenotomy) young and geriatric mice treated with PGDHi or vehicle (**Fig. 5a**). The snRNA-seq approach enabled comprehensive profiling of both myonuclear and non-myonuclear compartments. In total, ∼200,000 nuclei were sequenced across conditions. We identified and labeled 21 nuclei types, whose marker genes are shown in **Fig. 5b** by a dot plot, and we visualized them by UMAP (**Fig. 5c**). Our analysis was able to detect all major skeletal muscle populations, including distinct myonuclear subtypes (Myo_I, Myo_IIa, Myo_IIb, Myo_IIxb, Myo_New, NMJ, MTJ), as well as diverse muscle stem cell (MuSC) populations (MuSCs, Activated_MuSCs, Myocytes), fibro/adipogenic progenitors (FAPs), endothelial cells (Endothelial_Cell_I, Endothelial_Cell_II), smooth muscle cells (SMC), Schwann cells, adipocytes, and multiple immune populations (B_cells, T_cells, Dendritic_cells, and Monocyte/Macrophages) (**Fig. 5b, c**).

**Figure 5.**
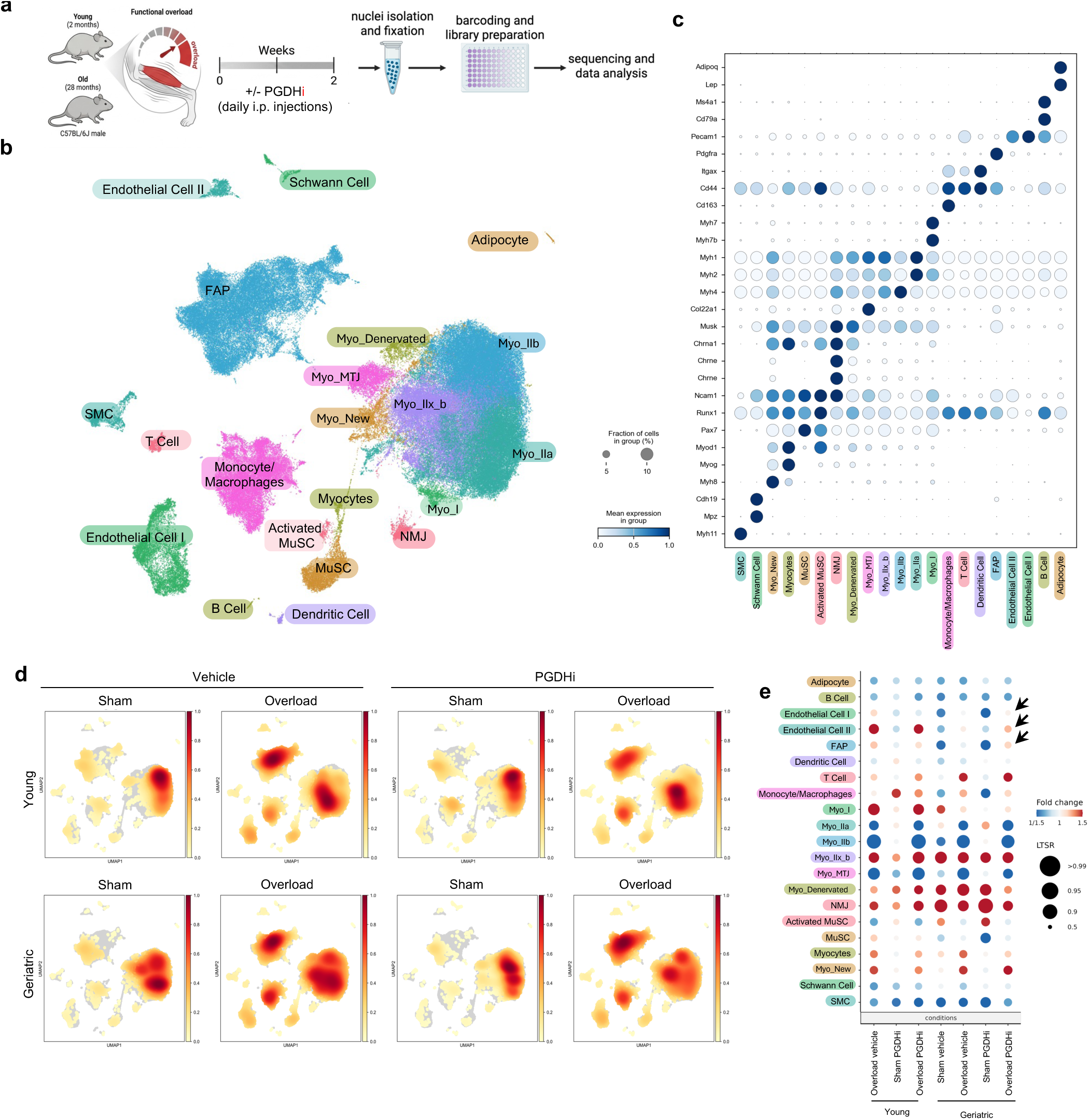
Single-nuclei RNA sequencing of overloaded and PGDHi-treated skeletal muscle. (a) Experimental design: young (2-month-old) and geriatric (28-month-old) mice underwent overload surgery and were treated with vehicle or PGDHi for 14 days. Plantaris muscles were collected for single-nuclei RNA sequencing. (b) Dot plot of marker gene expression used to identify cell types across all conditions. Dot size indicates the fraction of cells expressing each gene, and color intensity indicates mean expression level. (c) UMAP visualization of all nuclei, showing clustering of major skeletal muscle populations, including myonuclear subtypes (Myo_I, Myo_IIa, Myo_IIb, Myo_IIx_b, Myo_New, NMJ, MTJ, denervated), muscle stem cells (MuSC, activated MuSCs), myocytes, fibro/adipogenic progenitors (FAPs), endothelial cells, immune populations (B cells, dendritic cells, monocyte/macrophages, T cells), smooth muscle cells (SMCs), Schwann cells, and adipocytes. (d) Density plots of nuclei distribution across UMAP space for young and geriatric mice, in sham or overload conditions, with or without PGDHi. (e) Dot plot of changes in cell population abundance across conditions (compared to young sham vehicle), shown as log fold change (color) and LTSR score (dot size).

We hypothesized that cell composition would be altered in the course of plantaris muscle hypertrophy. Indeed, we observed a robust remodeling of muscle tissue and a shift in the proportions of diverse cell types in response to overload (**Fig. 5d, e**). In both young and aged mice, overload led to increased representation of MuSCs, activated MuSCs, FAPs, monocyte/macrophages, and endothelial cells. Importantly, in geriatric muscle, PGDHi treatment markedly enriched the FAPs and endothelial cells beyond the vehicle-treated condition, suggesting that 15-PGDH inhibition potentiates stromal and vascular responses during hypertrophy.

To resolve transcriptional programs altered by overload and PGDHi we performed differential expression analysis (PyDESeq2)^40,41^ comparing sham vehicle with overload vehicle or PGDHi vehicle groups, followed by GO enrichment. This analysis revealed that PGDHi treatment most strongly impacted the Myo_New and Myo_IIb myonuclear populations (**Fig. 6a**). In Myo_New, which are the newly incorporated myonuclei, PGDHi augmented expression of genes related to myogenic differentiation such as myogenin and myomaker (e.g., *Myog*, *Cd81*, *Mymk*) and muscle structural proteins including various myosin heavy chains (*Myh8*, *Myh3*, *Myh1*), with GO terms enriched for muscle development, contraction, and actin cytoskeleton organization (**Fig. 6b**). This aligns with our histological evidence of increased new myonuclear incorporation (Figure 4d). Notably, in Myo_IIb nuclei, PGDHi enhanced expression of genes linked to hypertrophy, IGF, and mTOR signaling (**Fig. 6c**), suggesting that type IIb myonuclei are central hubs of anabolic signaling in geriatric overloaded muscle when treated with PGDHi.

**Figure 6.**
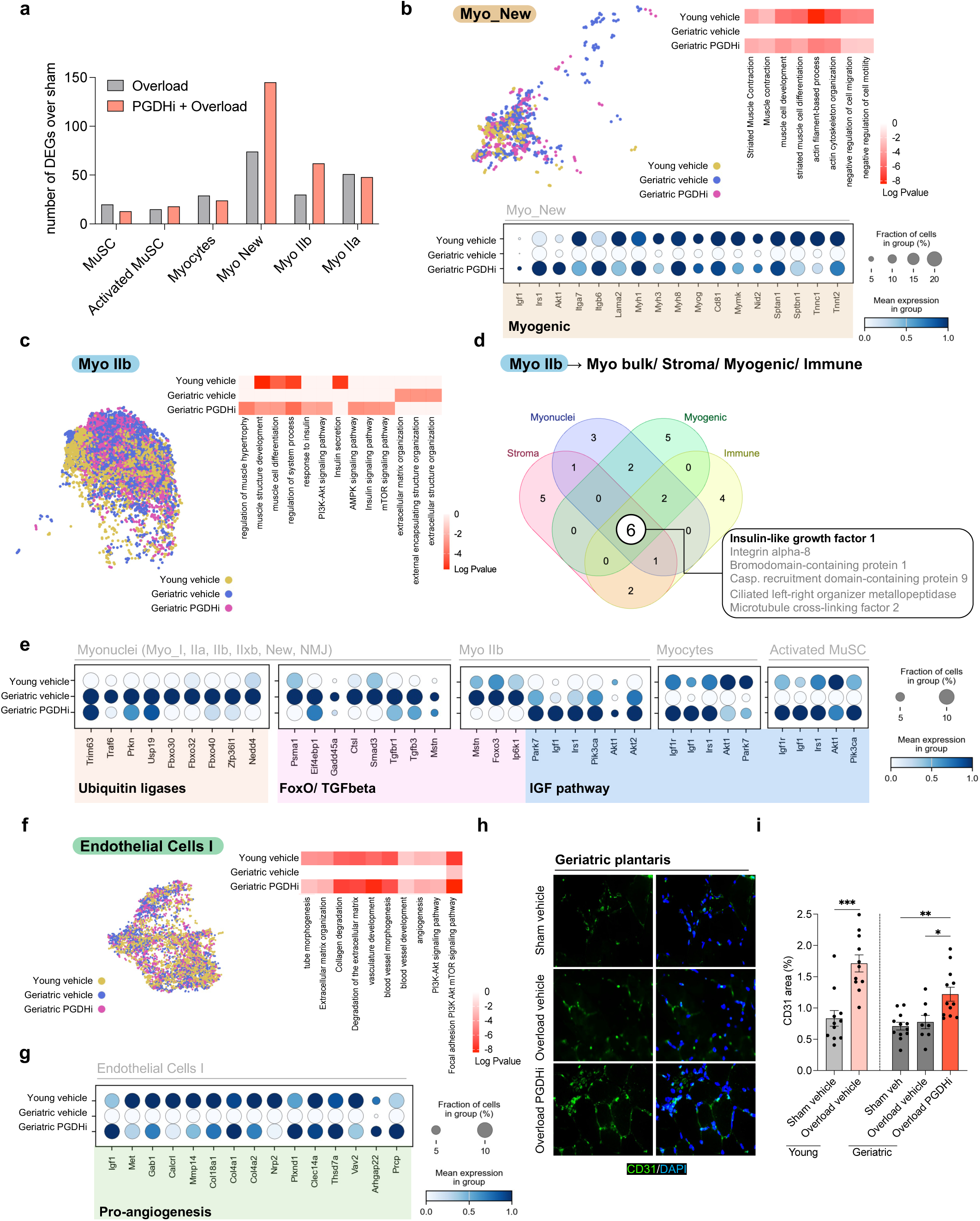
PGDHi enhances IGF signaling and vascular growth in overloaded aged muscle. (a) Bar graph showing the number of differentially expressed genes (DEGs) over sham in selected cell populations (MuSC, activated MuSC, myocytes, Myo_New, Myo_IIb, Myo_IIa) from overloaded or PGDHi-treated overloaded muscles. (b) Analysis of the Myo_New population. UMAP visualization of Myo_New nuclei from young vehicle, geriatric vehicle, and geriatric PGDHi groups (left). Gene ontology (GO) terms enriched among DEGs in Myo_New nuclei of overloaded muscles (right heatmap). Dot plot of representative DEGs in Myo_New nuclei across groups (bottom), with dot size indicating fraction of nuclei expressing each gene and color indicating mean expression. (c) UMAP visualization of Myo_IIb nuclei from young vehicle, geriatric vehicle, and geriatric PGDHi groups (left), with enriched gene ontology (GO) terms shown as a heatmap (right). (d) Venn diagram of predicted ligand–receptor interactions from Myo_IIb to bulk myonuclei, stromal, myogenic, and immune compartments, highlighting IGF1 as a shared interactor. (e) Dot plots showing expression of genes related to ubiquitin ligases, FoxO/TGFβ signaling, IGF pathway, and representative targets in myocytes and activated MuSCs across conditions. Dot size indicates the fraction of cells expressing the gene, and color intensity represents mean expression. (f) UMAP visualization of endothelial cell I populations (left) with enriched GO terms shown as a heatmap (right). (g) Dot plots of pro-angiogenic DEGs in endothelial cells across conditions. (h) Representative immunofluorescence images of CD31+ endothelial cells (green) in plantaris muscle sections from geriatric mice under sham, overloaded vehicle, and overloaded PGDHi conditions. Nuclei counterstained with DAPI (blue). Scale bar, 100 µm. (i) Quantification of CD31+ area as a percentage of total tissue area (n = 8-12 per group). Each dot represents one animal. Data represent mean ± SEM. \**P* < 0.05, \*\**P* < 0.01, \*\*\**P* < 0.001, **** *P* < 0.0001 by unpaired t-test or one-way ANOVA with post hoc test, as appropriate.

We therefore hypothesized that paracrine signaling from fiber type IIb might play a major role in the hypertrophic response seen with PGDHi treatment. To further assess potential communication between cell types, we used the Ligand–Receptor Inference Analysis (LIANA) pipeline^42,43^, grouping recipient cells into Stromal (FAPs, Endothelial I, Endothelial Cell II, and SMC), Myogenic (MuSCs, Activated_MuSCs, Myocytes), Myonuclei (Myo_I, Myo_IIa, Myo_IIb, Myo_IIxb, Myo_New, NMJ), and Immune cells (B_cells, T_cells, Dendritic_cells, and Monocyte/Macrophages). This analysis identified Myo_IIb myonuclei as a major source of insulin-like growth factor 1 (IGF1), a protein growth factor that is known to play a major anabolic role and predicted to signal broadly across all four cellular compartments, including stroma, immune, myogenic and myonuclear (**Fig. 6d, S5a–c**). Because IGF1 is a secreted growth factor, these data position type IIb myonuclei as a paracrine source rather than a cell-autonomous responder, with IIb-derived IGF1 signaling to neighboring cells and fibers—including the type IIa fibers whose cross-sectional area was preferentially increased by PGDHi (Fig. 3g)—to coordinate the hypertrophic response.

In conjunction with this anabolic effect, in the Myonuclei populations PGDHi caused a repression of multiple atrophy-associated genes, including classical ubiquitin ligases (*Trim63/MuRF1*, *Fbxo32/Atrogin-1*), *Foxo* transcription factors, and TGFβ pathway components. Moreover, PGDHi suppressed *Ip6k1*, a kinase recently implicated in promoting proteostasis defects and muscle atrophy^44^. In parallel, PGDHi enhanced activation of the well characterized growth promoting IGF1 pathway, with upregulation of *Igf1*, *Irs1*, *Akt1/2*, and *Rps6kb1* not only in Myo_IIb nuclei but also in myocytes and activated MuSCs—the progenitor populations that contribute directly to Myo_New incorporation (**Fig. 6e**). Together, these findings indicate that PGDHi simultaneously suppresses catabolic programs and activates pro-hypertrophic gene networks, shifting the balance of transcription of aged overloaded muscle tissue cells toward growth.

In addition to these myonuclear and myogenic progenitor effects, PGDHi also influenced the stromal and vascular compartments that were enriched in the cell composition analysis (**Fig. 5d, e**). Endothelial cell populations displayed PGDHi-dependent enrichment of pathways related to angiogenesis, including tube morphogenesis, extracellular matrix remodeling, and vascular development (**Fig. 6f, g**). Supporting these transcriptional signatures, CD31 immunostaining demonstrated that overload increased vascularization in young but not geriatric muscle, whereas PGDHi restored angiogenic responses in geriatric overloaded muscle (**Fig. 6h, i**). Thus, PGDHi not only enhances anabolic signaling in myonuclei and suppresses catabolic pathways, it also rejuvenates vascular remodeling, all of which converges to reinforce the hypertrophic response in aged skeletal muscle.

### Inhibition of IGF1 signaling blunts PGDHi-induced muscle growth and function

We hypothesized that a major mechanism by which PGDHi acts to augment muscle function is via IGF1. The IGF1 pathway has long been linked to skeletal muscle growth and is known to be dysregulated with aging, contributing to age-associated muscle weakness^45^. However, 15-PGDH and PGE2 have not previously been directly implicated in regulating IGF1. Our single-nuclei RNA-seq analysis revealed that PGDHi treatment upregulated the IGF1 signaling pathway at the transcriptional level in overloaded old muscle, and based on ligand-receptor (LIANA) analysis we found that *Igf1* expression localized predominantly to Myo_IIb myonuclei was predicted to target stromal, myogenic, myonuclear, and immune populations (**Fig. 6d**).

To confirm activation of the IGF1 signaling pathway by PGDHi at the protein level, we performed a phosphorylation array targeting IGF1R-related signaling components. Plantaris muscles from 28-month-old geriatric mice subjected to 6 days of overload and daily PGDHi or vehicle treatment were analyzed. We selected this day-6 time point to capture early signaling events during the active remodeling phase of overload, before the later hypertrophic endpoint, allowing us to assess pathway activation rather than secondary consequences of increased muscle size. PGDHi treatment increased phosphorylation of IGF1R (Tyr1161), IRS-2 (Ser731), Akt (Thr308, Ser473), p70S6K (Thr421/Ser424), CREB (Ser133), and FOXO1/3, consistent with activation of the IGF1–Akt–mTOR anabolic signaling cascade (**Fig. 7a**).

**Figure 7.**
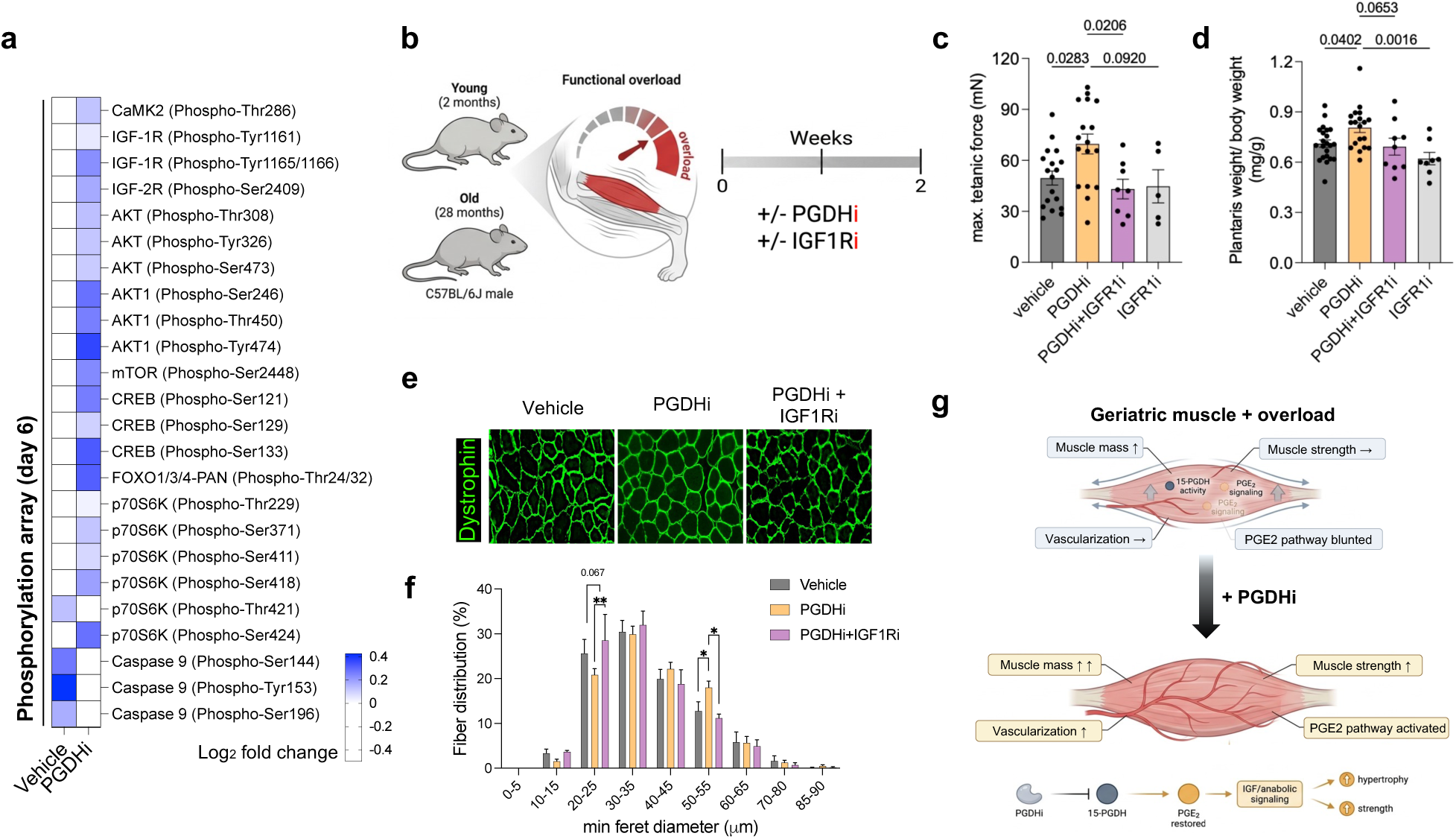
IGF1 signaling is required for the hypertrophic effects of PGDHi in aged muscle. (a) Phosphorylation array of IGF1R-related signaling proteins from overloaded plantaris muscles of geriatric mice treated with vehicle or PGDHi for 6 days. Heatmap shows log₂ fold change relative to vehicle. (b) Experimental schematic of young and aged mice subjected to overload and treated with PGDHi, with or without IGF1 receptor inhibitor (IGF1Ri), for two weeks. (c) Absolute plantarflexor force in geriatric mice across treatment groups (n = 5-18 per group). (d) Plantaris muscle weight in geriatric mice across treatment groups (n = 5-18 per group). (e) Representative immunofluorescence images of dystrophin-stained muscle cross-sections from geriatric mice treated with vehicle, PGDHi, or PGDHi + IGF1Ri. Scale bar, 100 µm. (f) Fiber size distribution of plantaris myofibers in geriatric mice treated as in (e). (g) Schematic summary illustrating impaired overload response in aged muscle (top), and how PGDHi restores hypertrophic adaptations through enhanced PGE2 bioavailability and IGF1 induction in type IIb fibers (bottom). Data represent mean ± SEM. \**P* < 0.05, \*\**P* < 0.01, \*\*\**P* < 0.001, **** *P* < 0.0001 by unpaired t-test or one-way ANOVA with post hoc test, as appropriate.

To directly test the requirement for IGF1 signaling in the hypertrophic effects of PGDHi, we performed overload experiments in 28-month-old geriatric mice treated with PGDHi, with or without an IGF1 receptor inhibitor (IGF1Ri) (**Fig. 7b**). In geriatric mice, PGDHi significantly increased plantar flexor force and muscle mass, whereas IGF1Ri blunted both effects (**Fig. 7c, d**). Histologic analysis of dystrophin-stained sections showed that PGDHi shifted fiber size distribution toward larger diameters, an effect attenuated by IGF1Ri (**Fig. 7e, f**). Together, these results indicate that PGDHi enhances muscle growth and function in geriatric overloaded muscle, at least in part, through an IGF1-dependent mechanism (**Fig. 7g**). These data place PGE2 upstream of IGF1 and show its crucial role in regulating muscle anabolic function.

## Discussion

Age-related loss of muscle mass and strength impairs mobility and increases morbidity and mortality in older adults^3–5^. A major barrier to maintaining and building muscle in later life is anabolic resistance, in which aged muscle fails to convert anabolic stimuli such as mechanical loading (i.e., resistance training) into sustained gains in muscle mass and strength^10,44,46^. We recently identified 15-PGDH, the PGE2-degrading enzyme, as a hallmark of aging that accumulates across multiple tissues and constrains PGE2 levels. Overexpressing 15-PGDH in young muscles reduced PGE2 levels and induced muscle fibers to shrink and weaken, similar to the atrophy and strength decline that accompanies aging. Conversely, 15-PGDH inhibition preserved muscle mass and strength in aged mice^25,47^. Our prior work identified PGE2 as a regulator of muscle stem cell proliferation and survival in the context of injury^23,34^ and as a regulator of neuromuscular synapses capable of restoring neuromuscular connectivity. Here we show that 15-PGDH inhibition can overcome anabolic resistance and that it does so by a previously unrecognized paracrine mechanism.

We found that impaired PGE2 signaling plays a central role in anabolic resistance in aged skeletal muscle. In young animals, mechanical overload robustly induced expression of COX-2, a key enzyme in the PGE2 biosynthetic pathway. In aged muscle, this induction was blunted and associated with diminished gains in hypertrophy and strength. Pharmacologic inhibition of 15-PGDH in overloaded muscles of geriatric mice increased PGE2 bioavailability, promoted protein synthesis, and rescued muscle growth and strength — findings consistent with a causal role for PGE2 insufficiency in anabolic resistance. These results provide mechanistic insights into previous findings showing that COX-dependent prostaglandin production rises with resistance-type stimuli and that short-term COX-2 inhibition blunts acute myofibrillar protein synthesis and hypertrophic signaling in humans and rodents^34,48–51^. Here we uncover an unexpected link between 15-PGDH and IGF1, and define the paracrine mechanism through which they drive muscle growth. We further show that just two weeks of 15-PGDH inhibition, combined with mechanical overload, produces robust gains in both muscle mass and strength in geriatric mice.

Beyond its established effects on muscle stem cells, 15-PGDH inhibition during overload produced two complementary actions in aged muscle. First, PGDHi restored an intercellular growth program that amplified overload adaptation in aged muscle. Our data linked PGE2 to the well-known mediator of anabolic growth, IGF1. Single-nuclei transcriptomics and ligand–receptor inference identified type IIb myonuclei as a source of IGF1-linked signaling induced by 15-PGDH inhibition, with predicted paracrine signals to stromal, myogenic, myonuclear, and immune cells — consistent with myofibers acting as active signaling hubs during loading rather than passive recipients of systemic cues^52–54^. IGF1R blockade eliminated the hypertrophic and functional benefits of 15-PGDH inhibition, demonstrating that IGF1 is required downstream of PGDHi for the full anabolic response. These findings identify PGE2 catabolism as a druggable node that operates upstream of the IGF1 signaling axis, capable of restoring anabolic growth in aged muscle.

Second, 15-PGDH inhibition dampened a broader catabolic, atrophy-promoting program that constrains muscle growth with aging. Myostatin (*Mstn*) is a well-established negative regulator of skeletal muscle mass and among the most intensively pursued therapeutic targets in muscle biology, with genetic and pharmacologic inhibition producing robust hypertrophy in multiple preclinical settings^55–59^PGDHi suppresses this anti-growth program at multiple nodes, downregulating TGF-β–linked restraint — including *Mstn* and *Tgfbr1* — together with FoxO-associated catabolic signaling, reflected by reduced expression of the canonical atrogenes MuRF1 (*Trim63*) and Atrogin-1 (*Fbxo32*). These atrogenes are elevated with age in rodents and humans^60,61^, and diverse interventions that improve age-associated muscle phenotypes — including rapalogs, sestrins, and apelin — converge on dampening FoxO-linked catabolic programs alongside improved anabolism^62–64^. In overloaded aged muscle, PGDHi restores a permissive pro-growth environment by coupling IGF1-dependent anabolic signaling with suppression of TGF-β/FoxO-linked catabolism.

Our findings speak to a broader theme: that PGE2 biology is pleiotropic and tissue-context dependent, and that by modulating PGE2 availability it is possible to orchestrate repair through distinct cellular routes. In articular cartilage, recent work from our group showed that 15-PGDH inhibition promotes cartilage regeneration primarily through changes in the gene expression program within resident chondrocytes—shifting populations away from hypertrophic-like programs toward matrix-synthesizing articular chondrocyte states—rather than through stem cell/progenitor expansion^65^. More generally, prostaglandin signaling—particularly via EP receptor–linked cAMP/PKA pathways—can oppose profibrotic TGF-β/Smad programs in other tissues such as lung^66,67^ and heart^68–70^, highlighting that downstream mechanisms are shaped by the responding cell types and their receptors. In skeletal muscle specifically, our data suggest that during overload, preventing PGE2 catabolism drives paracrine signaling that promotes secondary anabolic mediators (including IGF1) and suppresses atrophy-promoting factors.

In summary, the inability to build muscle in aging diminishes quality of life, even for those who exercise. Our results identify impaired PGE2 signaling as a critical driver of age-related deficits in muscle growth capacity. By inhibiting 15-PGDH, we restore PGE2 bioavailability, induce an IGF1-dependent paracrine program, and re-engage anabolic signaling to rescue hypertrophy and strength in aged skeletal muscle. These findings suggest that PGDHi acts by reprogramming a lipid-metabolite signal—raising PGE2 availability to re-engage anabolic growth—and may thereby function as an “exercise amplifier,” a pharmacological strategy that enhances the anabolic response to mechanical loading and complements resistance training and possibly other anabolic interventions in late life.

## Methods

### Animals and Study Design

All animal experiments were performed under procedures approved by the Stanford University Administrative Panel on Laboratory Animal Care (APLAC, protocol # IACUC-32909). Male C57BL/6J mice were obtained from the National Institute on Aging (NIA) aged rodent colony or from Jackson Laboratory (young controls). Mice were housed in a temperature-controlled facility (20–22 °C) with a 12-hour light/dark cycle and ad libitum access to food and water.

For overload-induced hypertrophy experiments, young (2 months) and geriatric (28 months) mice were randomized (based on body weight, and baseline plantar flexion force).

### Tenotomy (mechanical overload) model

Mechanical overload–induced hypertrophy was produced by Achilles tendon tenotomy to functionally unload the major plantar flexor synergists while leaving the plantaris tendon intact, as described previously^28^. Briefly, adult mice were anesthetized under aseptic conditions, and a small skin incision was made to expose the Achilles tendon complex. The Achilles tendon was transected to unload the gastrocnemius/soleus complex, thereby increasing mechanical load on the intact plantaris. The wound was closed with sutures, and animals received peri-operative analgesia and were monitored daily for recovery and wound healing.

### Tissue collection and time course

Overloaded plantaris muscles were harvested at multiple post-tenotomy time points spanning acute to remodeling phases, up to 14 days after surgery, as indicated for each experiment. At each time point, muscles were collected for downstream analyses including wet mass, histology/immunostaining, RNA/protein assays, mass spectrometry, and contralateral limbs and/or sham-operated animals were used as controls where applicable.

### Drugs preparation

SW033291 (PGDHi) (ApexBio cat #A8709) was dissolved in 10% ethanol, 5% Cremophor EL (Sigma-Aldrich C5135), and 85% D5W (5% dextrose in water) as previously described^47^ for intraperitoneal injection (5 mg/kg daily). Indomethacin (Sigma-Aldrich cat#I7378) was dissolved in 10% ethanol, 5% Cremophor EL (Sigma-Aldrich C5135), and 85% D5W (5% dextrose in water) for intraperitoneal injection (2.5 mg/kg daily). IGF1 receptor inhibitor (Novus Biologicals cat# NBP1-77680PEP) was prepared in DMSO and diluted in PBS and administered intraperitoneally at 50 mg/kg daily.

### Force measurements

Isometric torque of the ankle plantar flexors was measured in anesthetized mice (3% isoflurane in oxygen) using a servomotor system (Aurora Scientific, Model 300C-LR). The foot was secured to a footplate, and percutaneous platinum–iridium electrodes were placed near the tibial nerve. The ankle joint was held at 90°, and tetanic contractions were evoked by 0.1 ms square pulses at 150 Hz. Three contractions were performed per mouse with 1-min rest intervals. For endpoint overload experiments, tetanic and specific force were determined by normalizing maximum force to plantaris mass.

### Protein extraction and Western Blot

Plantaris muscles were flash-frozen in liquid nitrogen and homogenized in RIPA buffer with protease and phosphatase inhibitors. Protein concentration was determined by BCA assay, and equal amounts were resolved by SDS-PAGE and transferred to PVDF membranes. Membranes were blocked in 5% milk and probed with antibodie against 15-PGDH. Ponceau was used to normalize to protein content. Blots were imaged using chemiluminescence and quantified with ImageJ.

### Phosphorylation array

Activation of IGF1R-related signaling pathways was assessed using the IGF1R Phospho Antibody Array (Full Moon BioSystems, Cat. #PTK124), according to the manufacturer’s protocol. Plantaris muscles from 28-month-old mice subjected to 6 days of overload and treated with PGDHi or vehicle were homogenized in the provided lysis buffer. Equal amounts of protein from three biological replicates per group were pooled to generate sufficient input for array analysis. Following labeling with biotin, samples were hybridized to the array slides containing capture antibodies for IGF1R pathway targets. After incubation with Cy3-streptavidin, fluorescence intensities were scanned with an Agilent DNA microarray scanner and quantified using GenePix Pro software (Molecular Devices). Signal intensities were normalized to internal controls and compared between treatment groups.

### Real Time RT-qPCR

Total RNA was extracted from frozen plantaris muscles using the RNeasy Fibrous Tissue Mini Kit (Qiagen, Cat. #74704), optimized for fibrous tissues such as skeletal muscle. RNA concentration and purity were determined using a NanoDrop spectrophotometer (Thermo Fisher). For cDNA synthesis, 500 ng–1 μg of RNA was reverse-transcribed with the ReverTra Ace qPCR RT Master Mix (Toyobo, Cat. #FSQ-201) following the manufacturer’s protocol.

Quantitative PCR was performed using SYBR Green Master Mix (Bio-Rad) on a QuantStudio 6 Flex Real-Time PCR System (Applied Biosystems). Target genes included markers of muscle stem cell activation and differentiation (Pax7, Myf5, Myog, Mymx, Mymk), anabolic signaling (Igf1), and PGE2 biosynthesis/degradation (Cox2, 15-Pgdh, Ptger4). Relative gene expression was calculated using the ΔΔCt method with 18S rRNA (S18) as the internal control. All reactions were performed in triplicate, and melt-curve analysis confirmed amplification specificity. Primer sequences are listed in Table S1.

### Immunohistochemistry and EdU labeling

Frozen cross-sections (10 µm) were fixed in 4% paraformaldehyde (PFA) for 10 min at room temperature, followed by three washes in phosphate-buffered saline (PBS). Sections were blocked in Blocking One (Nacalai Tesque) and incubated overnight at 4 °C with primary antibodies against Laminin, Pax7, Dystrophin, and CD31, or with the Click-iT EdU detection kit (Invitrogen) following the manufacturer’s instructions. The next day, samples were washed three times for 10 min each with PBS containing 0.1% Tween-20 (PBS-T) and incubated for 1 h at room temperature with appropriate secondary antibodies. After three additional washes in PBS-T, nuclei were counterstained with DAPI and sections mounted with antifade mounting medium. Images were acquired using a Keyence BZ-X810 fluorescence microscope.

Fiber cross-sectional areas and Feret diameters were quantified in BZ-X800 software (Keyence). For proliferation assays, EdU was administered in drinking water (0.2 mg/mL) for 7 days. EdU+ and Pax7+ cells were quantified, and EdU+ myonuclei incorporated into myofibers were scored as new myonuclear accretion.

### Single nuclei RNA-sequencing

#### Nuclei isolation and library preparation

Single-nuclei suspensions were prepared from ∼30–50 mg of snap-frozen plantaris muscles from young and geriatric mice, following a modified version of the 10x Genomics “Nuclei Isolation from Complex Tissues” protocol. Briefly, tissue was minced in NP-40 lysis buffer (10 mM Tris-HCl pH 7.4, 10 mM NaCl, 3 mM MgCl₂, 0.1% Igepal, 1 mM DTT, 1 U/µL RNase inhibitor) until homogeneous, followed by gentle Dounce homogenization and lysis on ice. The suspension was filtered through a 40 µm strainer, washed in nuclei FACS buffer (1% BSA, 0.2 U/µL RNase inhibitor in PBS), and stained with 7AAD (Miltenyi). Nuclei were pelleted at 300 g for 10 min at 4 °C and resuspended in FACS buffer. Viable (7AAD⁻) nuclei were isolated by FACS (Sony SH800) and immediately fixed using the Evercode Cell Fixation v1 kit (Parse Biosciences), then stored at –80 °C for a maximum of two weeks before processing. Libraries were prepared using the Evercode WT Mega v1 kit (Parse Biosciences) according to the manufacturer’s instructions.

#### Sequencing

Libraries were quantified with a Qubit fluorometer (Thermo Fisher), and fragment size distribution was verified with an Agilent Bioanalyzer. Sequencing was performed on an Illumina NovaSeq 6000 with paired-end 150 bp reads, targeting ∼25,000 reads per nucleus.

#### Pre-processing and quality control

Raw sequencing data were processed using the Parse Biosciences pipeline to demultiplex, assign barcodes, align reads to the mouse reference genome (mm10), and generate nuclei count matrices. Subsequent analyses were conducted in Scanpy (v1.9.1). Low-quality nuclei (<100 detected genes), genes expressed in fewer than 5 nuclei, and nuclei with >5% mitochondrial reads were removed. Doublets were identified with Solo and excluded if the predicted doublet probability exceeded 90%. After filtering, a total of ∼200,000 high-quality nuclei across all experimental conditions were retained for downstream analysis.

#### Dimensionality reduction and clustering

Dimensionality reduction and batch correction were performed using scVI (via scvi-tools; n_latent = 30, n_layers = 3, n_hidden = 128, max_epochs = 100), correcting for sample and percent mitochondrial content. Uniform Manifold Approximation and Projection (UMAP, n_neighbors = 15) was used for visualization. Clusters were identified with the Leiden algorithm (resolution = 3) and assigned to known cell types based on canonical marker gene expression.

#### Differential gene expression and pathway analysis

Differential expression was performed by pseudobulking nuclei by sample and cell type, followed by analysis with pyDESeq2^41^. Genes with log₂ fold change >0.25 and adjusted p < 0.05 were considered differentially expressed. Gene ontology enrichment of DEG sets was carried out using Metascape^71^, with terms considered enriched at q < 0.1.

#### Ligand–receptor inference

Cell-cell communication was inferred using the Python package Liana^43^, which we adapted with several custom modifications suited to our analysis. Following the standard Liana workflow, we identified enriched ligand-receptor (L-R) interactions for every sender-receiver cell pair within each of the groups and recorded the corresponding total count of L-R interactions per sender-receiver pair in each group. To account for differences in overall interaction abundance, the L-R interaction count for a given sender-receiver pair was scaled relative to the sum of all L-R interactions detected across every sender-receiver pair within the same group. Using these normalized values, we then derived the change in L-R interaction abundance for each sender-receiver pair across the pairwise group comparisons.

We next assessed the biological pathways associated with the detected L-R interactions. A gene ontology (GO) pathway was treated as enriched by a particular L-R interaction only when both the ligand and the receptor belonged to that pathway. For each sender-receiver pair within a group, we tallied how frequently each pathway was enriched across the L-R interactions linking those two cell types. These tallies were normalized against the total number of pathways identified for the respective group. The resulting normalized enrichment values were then contrasted between groups (for each sender-receiver pair, and GO pathways were ordered according to the magnitude of change—whether increased or decreased—in each comparison. For each comparison, the ten most differentially enriched GO pathways per sender-receiver pair were presented as heatmaps, with red and blue indicating greater or lesser pathway enrichment, respectively, for a given sender-receiver pair between the two groups. All heatmaps depicting up- and downregulated features were produced in Python with the matplotlib and seaborn libraries.

### Statistics

Data are expressed as mean ± SEM. For two-group comparisons, two-tailed unpaired Student’s t-tests were used. Multi-group analyses were performed with one- or two-way ANOVA and Sidak or Tukey post hoc testing.

### LC-MS/MS Quantification of PGE2

PGE2 levels were measured as previously described^47^. Briefly, muscle tissue (Plantaris muscles) was harvested, weighted, snap-frozen in liquid nitrogen, homogenized in acetone/water (1:1 v/v) buffer containing butylated hydroxytoluene (BHT,0.005%), and spiked with deuterated internal standards (PGE2-D4). After protein precipitation, samples underwent a two-step liquid–liquid extraction with hexane and chloroform to enhance LC-MS/MS sensitivity, followed by evaporation under nitrogen and reconstitution in acetonitrile/0.1% acetic acid (3:7 v/v). All analyses were carried out by negative electrospray ionization and chromatographic separation on a UPLC BEH C18 column. LC-MS/MS analyses were performed on either a Shimadzu 20AD_xr_ Prominence LC system with Shimadzu 8030 triple quadrupole mass spectrometer or Waters Acquity I-class LC system with Waters Xevo TQ-XS triple quadrupole mass spectrometer. Calibration curves were prepared fresh with each run using authentic prostaglandin standards across a dynamic range of 0.025–500 ng/mL (200µL aliquot), with linearity confirmed (R^2^ > 0.99). Selected reaction monitoring (SRM) transitions (quantifier and qualifier ions) were selected for each analyte to ensure specificity. Quantification was based on the ratio of analyte to internal standard peak area, with calibration curve performed using a 1/X² weighting factor.

### In vivo protein synthesis

In vivo protein synthesis was quantified by measuring incorporation of [^35^S]-methionine into muscle proteins 4 days after Achilles tenotomy induced overload. Mice were fasted for 1 hr and then received an intraperitoneal injection of [^35^S]-methionine (5 μCi/g body weight; PerkinElmer) diluted in sterile saline. 30 min after tracer administration, plantaris muscles were dissected, weighed, and immediately frozen.

Frozen muscles were homogenized on ice in radioimmunoprecipitation assay (RIPA) buffer supplemented with a protease inhibitor cocktail (Roche) using a handheld homogenizer, and lysates were clarified by centrifugation at 4°C to remove debris. Proteins were precipitated with trichloroacetic acid (TCA), and the radioactivity associated with the protein pellet was measured by liquid scintillation counting. [^35^S]-methionine incorporation was normalized to tissue weight to obtain an index of in vivo protein synthesis.

## Acknowledgments

This work utilized the Xevo TQ-XS mass spectrometer system (RRID:SCR_018510) that was purchased with funding from National Institutes of Health Shared Instrumentation Grant S10OD026962.

## Funding

This work was supported by the Baxter Foundation, the Li Ka Shing Foundation, the Milky Way Research Foundation (grant 216064 to H.M.B.), the American Federation for Aging Research (grant 305693), NIH funding (R01AG02096115, R01AG069858, and RHG009674A to H.M.B.), and a Stanford Cardiovascular Institute Seed Grant awarded to H.M.B. and M.N. I.K. received support through the Stanford Bio-X Summer Undergraduate Research Program.

## Competing interests

H.M.B. is listed as an inventor on patents involving the use of PGE2 in muscle regeneration and rejuvenation, which have been licensed to Epirium Bio. In addition, H.M.B. acts as a consultant for Epirium Bio and serves on the company’s Scientific Advisory Board. K.J.S. is a co-founder and equity holder of Merrifield Therapeutics.

K.J.S. has received research funds from Merck, Eli Lilly, and Pfizer, and consulting fees from AbbVie and Novo Nordisk.

## Author contributions

M.N, and H.M.B. conceived the project and designed the study. M.N performed most of the experiments and led the analysis. Y.K.L., E.L.M., M.T., P.K., J.B., Z.Z., K.S., K.J., and I.K. carried out the mouse studies. M.N. and E.M. performed transcriptomic analyses. L.A. performed LC-MS/MS metabolite analyses. M.N., and H.M.B interpreted the data with input from all authors. M.N. and H.M.B wrote the first draft of the manuscript. All authors contributed to editing and revising the manuscript and approved the final version of the manuscript.

## Data and materials availability

All data supporting the findings of this study are available within the paper and its Supplementary Information files. The single-nuclei RNA-seq datasets will be deposited in GEO prior to publication. All other materials are available from the corresponding author upon reasonable request.

## Declaration of generative AI and AI-assisted technologies in the manuscript preparation process

During the preparation of this work, the authors used OpenAI’s ChatGPT solely to assist with language editing and improve clarity. The authors reviewed and edited the output as needed and take full responsibility for the content of the published article.

