## Supplemental Figures for "Restoration of Capacity to Build Muscle Strength in Geriatric Mice by Inhibition of the Gerozyme 15-Prostaglandin Dehydrogenase"

**Figure 1**

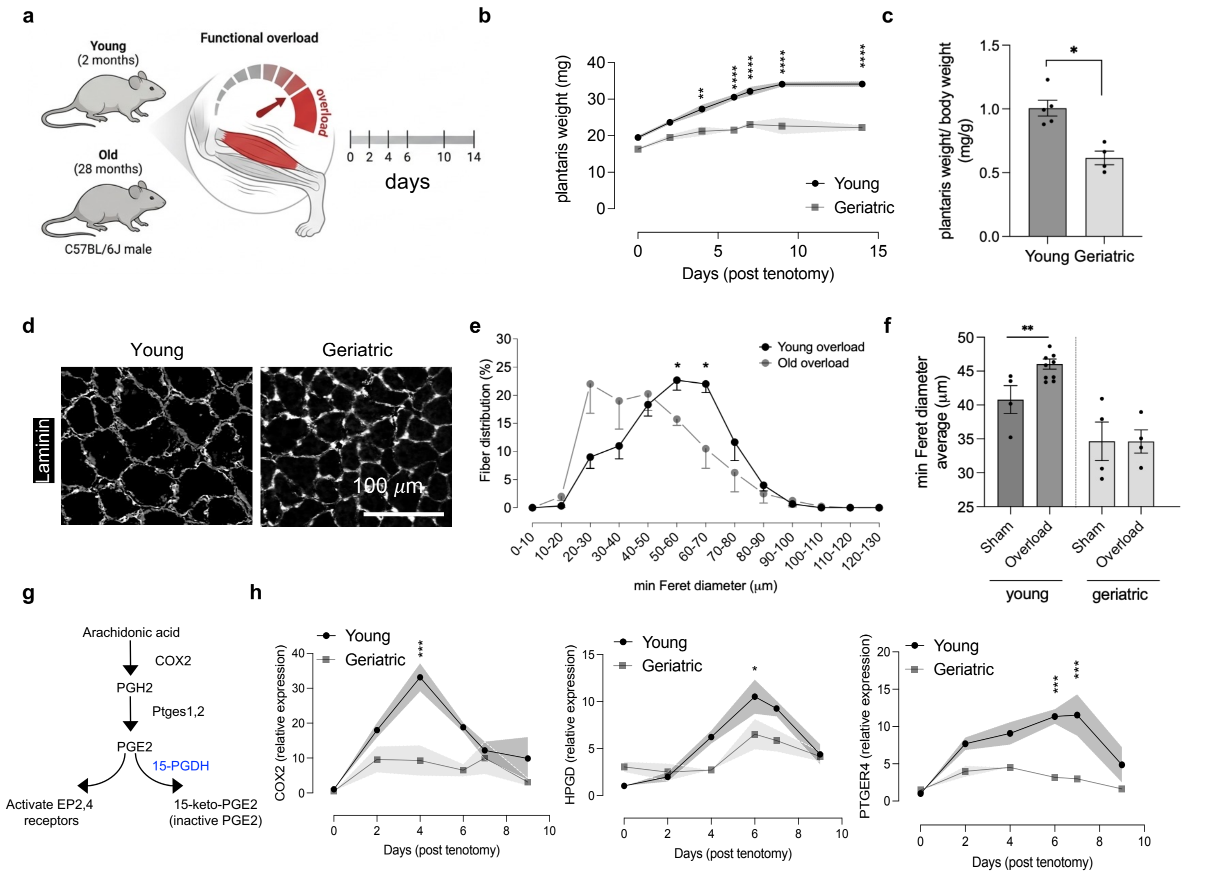


**Figure 1. Age-associated decline in overload-induced muscle hypertrophy and PGE2 signaling.**
(a) Schematic of experimental design in young (2-month-old) and geriatric (28-month-old) C57BL/6 male mice subjected to synergist ablation–induced overload.
(b) Time course of plantaris muscle weight following overload (n = 4 per group).
(c) Plantaris weight normalized to contralateral sham (n = 4 per group).
(d) Representative laminin-stained plantaris cross-sections from young and geriatric mice. Scale bar, 100 μm.
(e) Fiber size distribution curves of plantaris myofibers following overload.
(f) Quantification of minimal Feret diameters in sham and overloaded plantaris muscles.
(g) Schematic of the prostaglandin E2 biosynthetic and degradative pathway.
(h) Expression of *Cox2*, *Hpgd* (15-PGDH), and *Ptger4* measured by RT–qPCR at indicated days post-overload (n = 4 per group).
In (c) and (f), each dot represents one animal. In (b), (e), and (h), statistical significance indicates comparisons between geriatric and young animals at each time point. Data represent mean ± SEM. *$P < 0.05$, **$P < 0.01$, ***$P < 0.001$, **** $P < 0.0001$ by two-way ANOVA with post hoc test or unpaired *t*-test.

### Figure S1

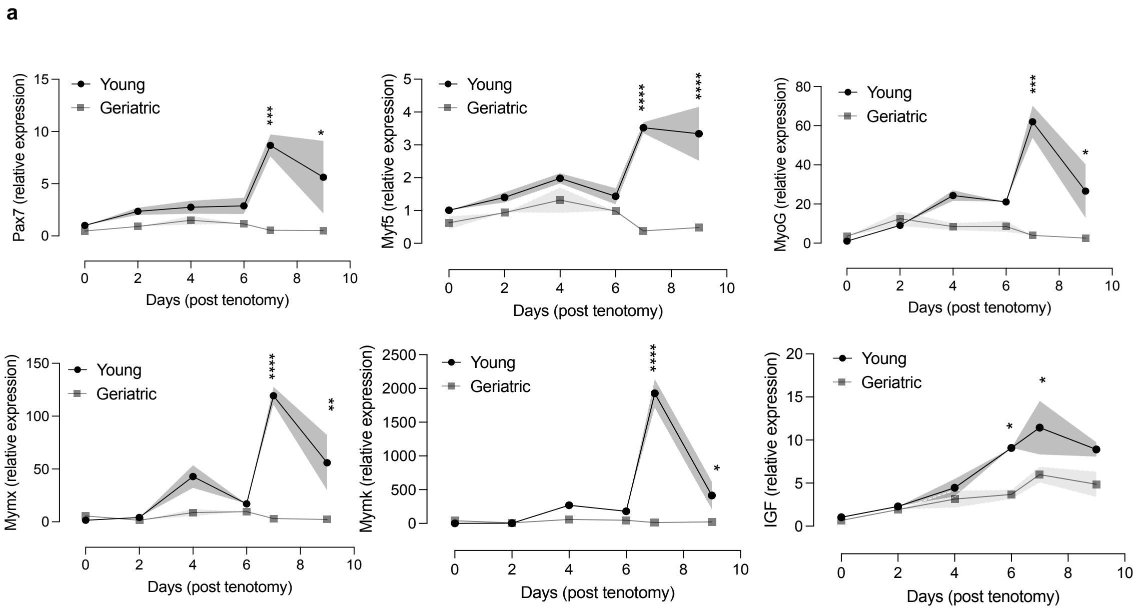


**Figure S1. Expression of myogenic genes during overload.**
(a) RT–qPCR analysis of *Pax7*, *Myf5*, *Myog*, *Mymx*, *Mymk*, and *Igf1* expression in plantaris muscles from young and geriatric mice at indicated days following overload (n = 4 per group).
Statistical significance indicates comparisons between geriatric and young animals at each time point. Data represent mean ± SEM. **P* < 0.05, ***P* < 0.01, ****P* < 0.001, **** *P* < 0.0001 by two-way ANOVA.

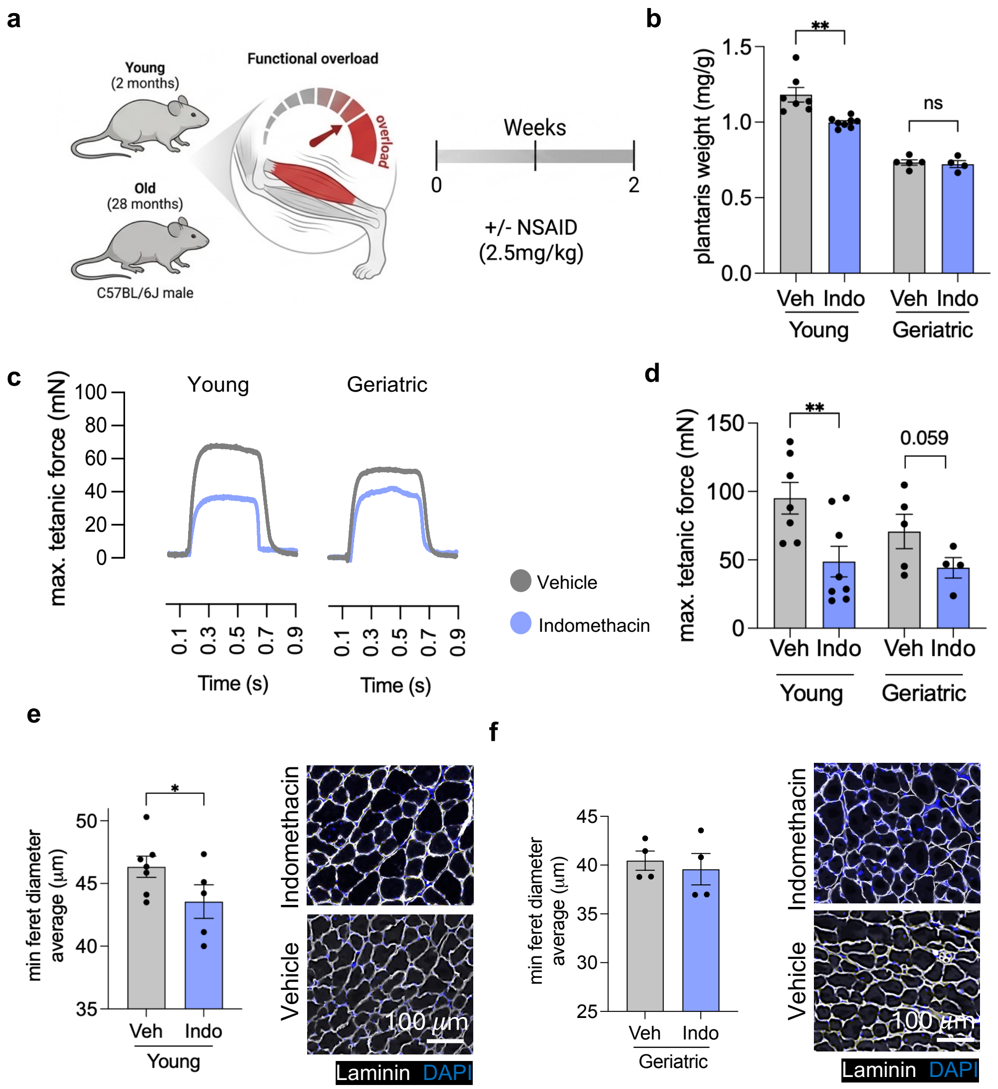


**Figure 2. Effect of COX inhibition on overload-induced hypertrophy in young and geriatric mice.**
(a) Experimental schematic: young (2-month-old) and geriatric (28-month-old) mice subjected to overload with daily intraperitoneal injection of vehicle or indomethacin (2.5 mg/kg).
(b) Plantaris muscle weight after 2 weeks of overload (n = 4-7 per group).
(c) Representative traces of tetanic force measurements in young and geriatric mice treated with vehicle or indomethacin.
(d) Quantification of maximal tetanic force (n = 4-7 per group).
(e) Minimal Feret diameter and representative laminin-stained cross-sections of plantaris muscle from young mice treated with vehicle or indomethacin. Scale bar, 100 µm.
(f) Minimal Feret diameter and representative laminin-stained cross-sections of plantaris muscle from geriatric mice treated with vehicle or indomethacin. Scale bar, 100 µm.
In (b), (d), (e) and (f) each dot represents one animal. Data represent mean ± SEM. Exact *P* values are shown, determined by unpaired *t*-test.

**Figure S2**

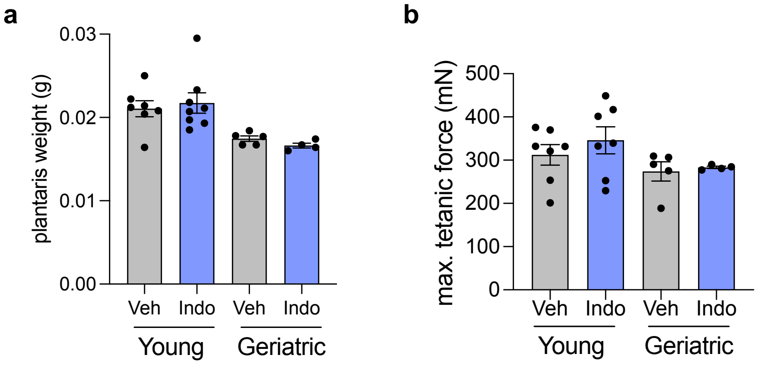


**Figure S2. Effect of indomethacin treatment on sham-operated muscles.**
(a) Plantaris muscle weight in young and geriatric mice treated with vehicle or indomethacin (n = 4-7 per group).
(b) Maximal tetanic force in sham-operated plantaris muscles from young and geriatric mice treated with vehicle or indomethacin (n = 4-7 per group).
Each dot represents one animal. Data represent mean ± SEM. Significance was determined by unpaired *t*-test.

### Figure 3

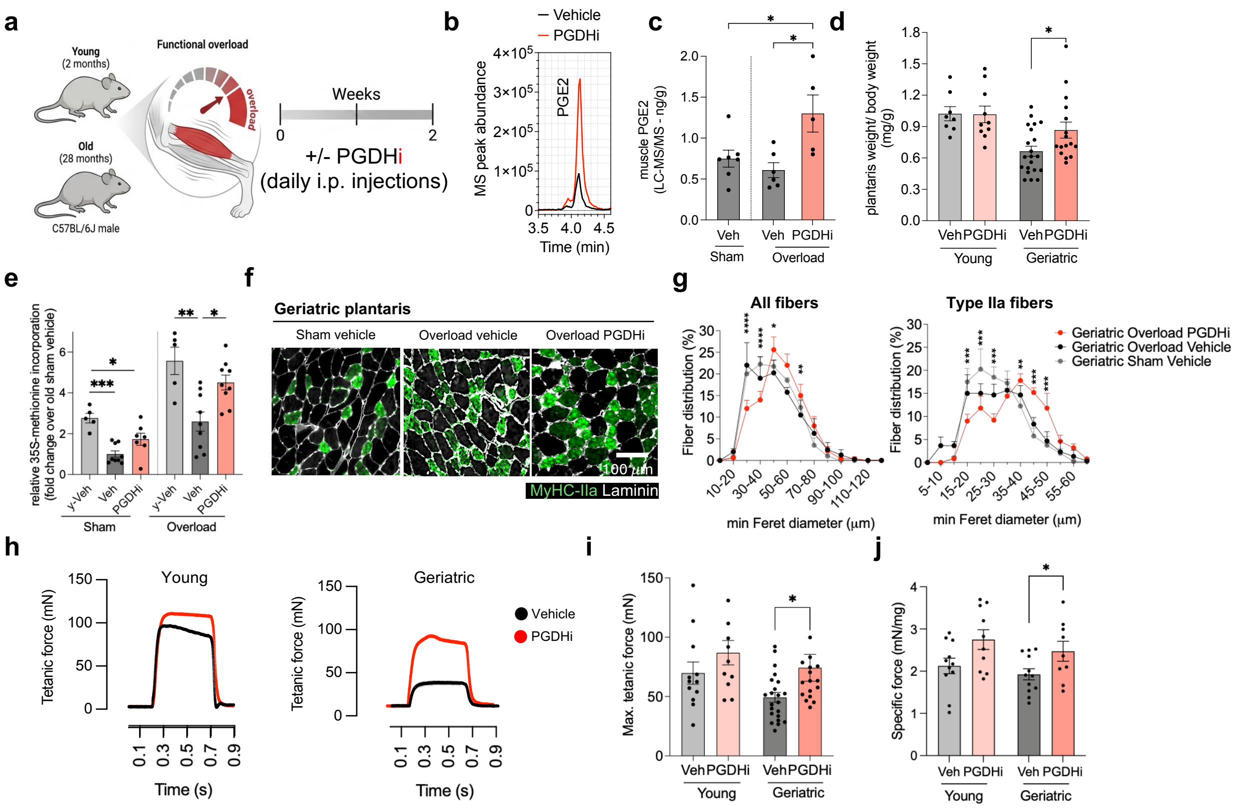


**Figure 3. 15-PGDH inhibition enhances hypertrophy and function in overloaded geriatric muscle.**
(a) Experimental schematic of overload-induced hypertrophy in young and geriatric mice treated with vehicle or 15-PGDH inhibitor (PGDHi) for 2 weeks.
(b) Representative LC–MS/MS chromatograms of PGE2 levels in overloaded geriatric plantaris muscles treated with vehicle or PGDHi. (c) Muscle PGE2 levels measured by LC–MS/MS in sham and overloaded plantaris muscles (n = 5-7 per group).
(d) Plantaris weight normalized to body weight in young and geriatric mice after overload with vehicle or PGDHi treatment (n = 8-21 per group).
(e) Levels of protein synthesis measured by [$^{35}$S]-methionine incorporation in plantaris muscles overloaded for four days.
(f) Representative laminin (white) and type IIa fiber (MyHC-IIa, green) immunostaining of geriatric plantaris muscle sections from sham and overloaded mice treated with vehicle or PGDHi. Scale bar, 100 µm.
(g) (left) Fiber size distribution of all fibers in geriatric overloaded muscles treated with vehicle or PGDHi (n = 4-8 per group). (right) Fiber size distribution of type IIa fibers in geriatric overloaded muscles treated with vehicle or PGDHi (n = 4-8 per group).
(h) Representative force traces of plantar flexor tetanic contractions from young and geriatric mice treated with vehicle or PGDHi.
(i) Maximal tetanic force of plantaris muscles in young and geriatric mice with vehicle or PGDHi treatment (n = 8-21 per group).
(j) Specific force (normalized to muscle weight) in young and geriatric mice treated with vehicle or PGDHi (n = 8-21 per group).
In (c), (d), (e), (i) and (j) each dot represents one animal. Data represent mean ± SEM. **P* < 0.05, ***P* < 0.01, ****P* < 0.001, **** *P* < 0.0001 by unpaired *t*-test.

#### Figure S3

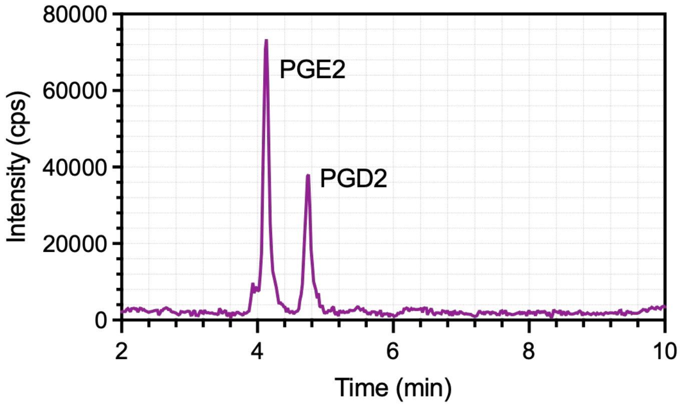


**Figure S3. Representative LC-MS/MS chromatogram of a standard mix showing chromatographic separation of PGE2 and PGD2.** Analyte peak intensities are expressed as cps, counts per second. LC-MS/MS data and calibration curves were acquired and analyzed using the LCMS-8030 triple quadrupole mass spectrometer (Shimadzu).

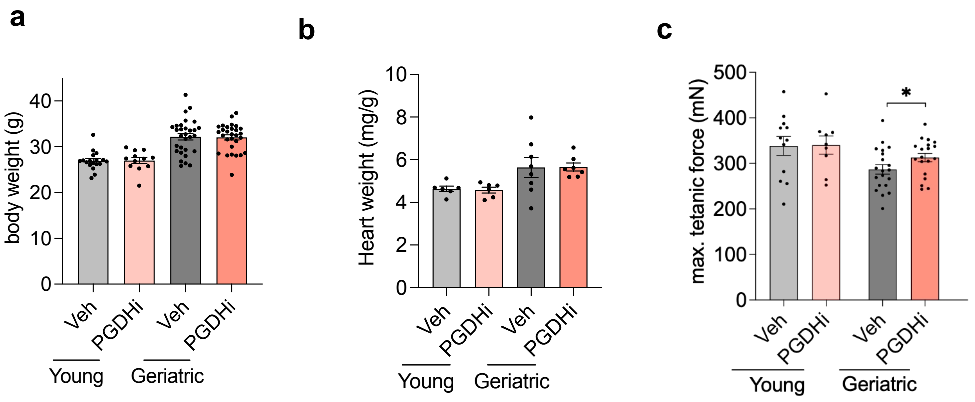


**Figure S4. No effect on body weight and heart weight of PGDHi treatment in young and geriatric mice.**
(a) Body weight of young and geriatric mice treated with vehicle or PGDHi (n = 8-21 per group).
(b) Heart weight normalized to body weight in young and geriatric mice treated with vehicle or PGDHi (n = 6-8 per group).
(c) Maximal tetanic force of the sham legs 2 weeks after treatment.
In (a), (b) and (c) each dot represents one animal. Data represent mean ± SEM. Significance was established by unpaired *t*-test.

### Figure 4

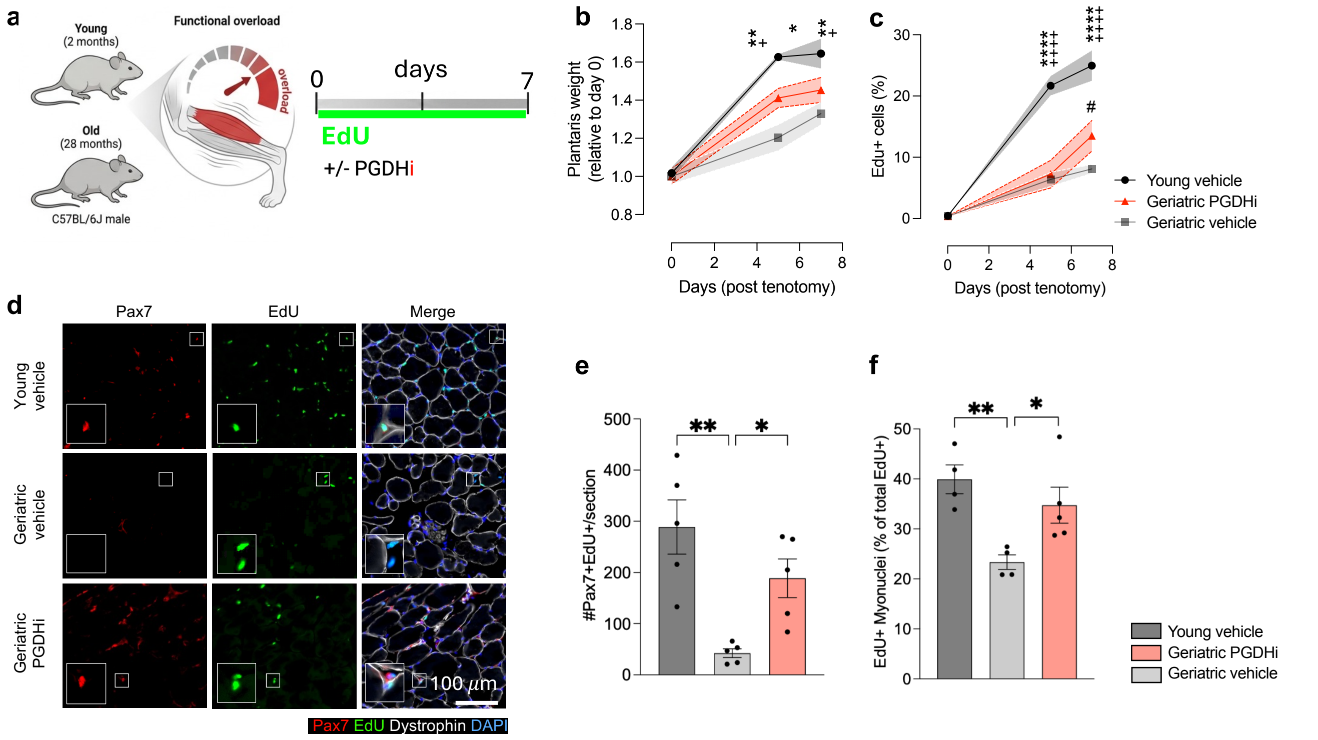


**Figure 4. PGDHi enhances muscle stem cell activity and myonuclear accretion in overloaded geriatric muscles**

(a) Experimental design: young (2-month-old) and geriatric (28-month-old) mice underwent overload surgery and were treated with vehicle or PGDHi for 7 days, with EdU administered continuously in drinking water.

(b) Plantaris muscle weight relative to baseline (n = 4-5 per group).

(c) Percentage of $EdU^{+}$ cells over the time course (n = 4-5 per group).

(d) Representative immunostaining of Pax7 (red), EdU (green), dystrophin (white), and DAPI (blue) in plantaris muscle sections. Insets highlight $Pax7^{+}EdU^{+}$ cells.

(e) Number of $Pax7^{+}EdU^{+}$ nuclei per section on day 7 after tenotomy (n = 4-5 per group).

(f) Number of $EdU^{+}$ myonuclei per section on day 7 after tenotomy (n = 4-5 per group).

In (e) and (f) each dot represents one animal. Data represent mean ± SEM. *$P < 0.05$, **$P < 0.01$, ***$P < 0.001$, **** $P < 0.0001$ by one-way ore two-way ANOVA with post hoc test.

### Figure 5

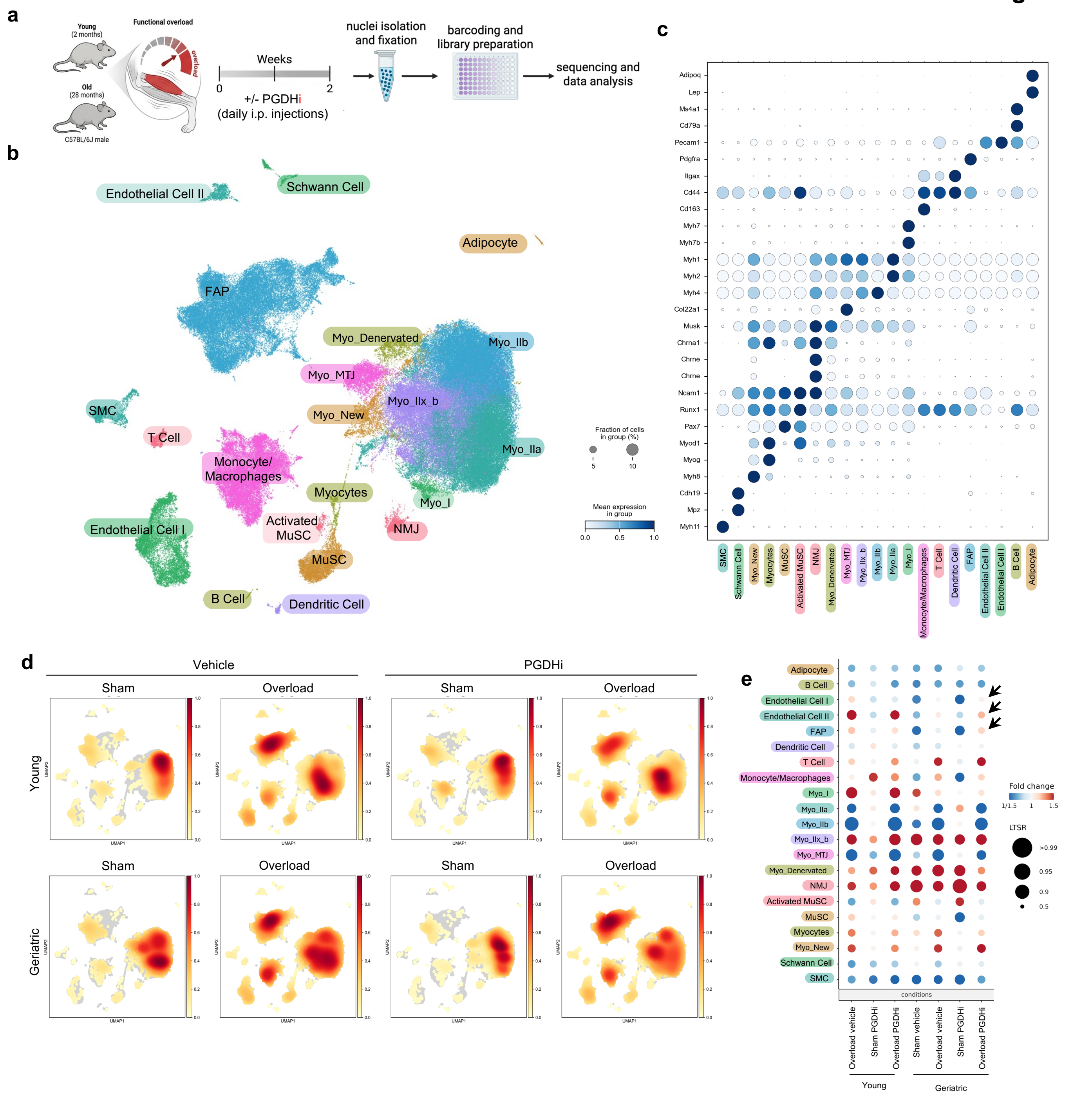


**Figure 5. Single-nuclei RNA sequencing of overloaded and PGDHi-treated skeletal muscle.**
(a) Experimental design: young (2-month-old) and geriatric (28-month-old) mice underwent overload surgery and were treated with vehicle or PGDHi for 14 days. Plantaris muscles were collected for single-nuclei RNA sequencing.
(b) Dot plot of marker gene expression used to identify cell types across all conditions. Dot size indicates the fraction of cells expressing each gene, and color intensity indicates mean expression level.
(c) UMAP visualization of all nuclei, showing clustering of major skeletal muscle populations, including myonuclear subtypes (Myo_I, Myo_IIa, Myo_IIb, Myo_IIx_b, Myo_New, NMJ, MTJ, denervated), muscle stem cells (MuSC, activated MuSCs), myocytes, fibro/adipogenic progenitors (FAPs), endothelial cells, immune populations (B cells, dendritic cells, monocyte/macrophages, T cells), smooth muscle cells (SMCs), Schwann cells, and adipocytes.
(d) Density plots of nuclei distribution across UMAP space for young and geriatric mice, in sham or overload conditions, with or without PGDHi.
(e) Dot plot of changes in cell population abundance across conditions (compared to young sham vehicle), shown as log fold change (color) and LTSR score (dot size).

### Figure 6

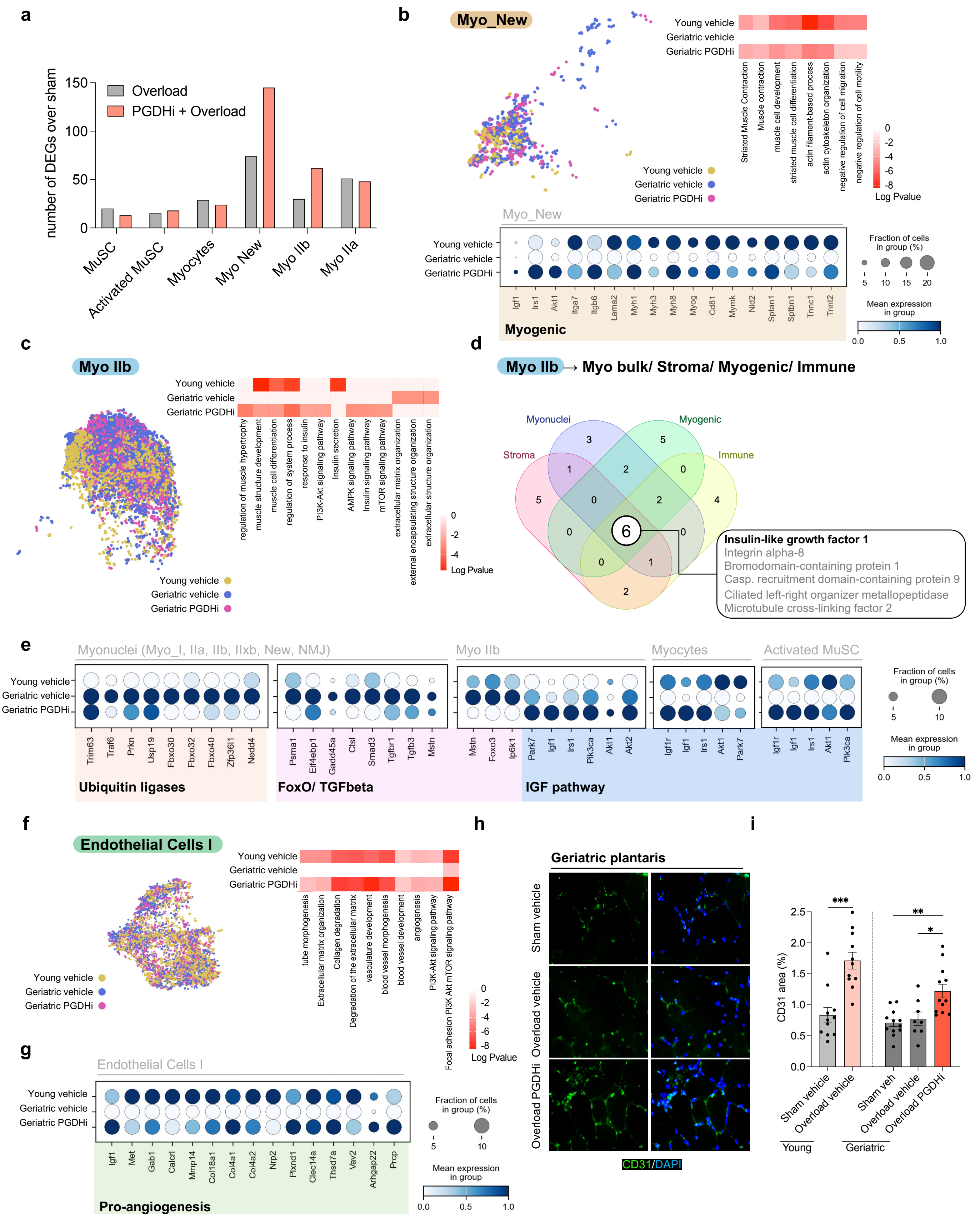
a
number of DEGs over sham
Overload
PGDHi + Overload
MuSC
Activated MuSC
Myocytes
Myo New
Myo IIb
Myo IIa
b
Myo_New
Young vehicle
Geriatric vehicle
Geriatric PGDHi
Striated Muscle Contraction
Muscle contraction
muscle cell development
striated muscle cell differentiation
actin filament-based process
actin cytoskeleton organization
negative regulation of cell migration
negative regulation of cell motility
Log Pvalue
Fraction of cells in group (%)
Mean expression in group
Igf1
Irs1
Akt1
Itga7
Itgb6
Lama2
Myh1
Myh3
Myh8
Myog
Cd81
Mymk
Nid2
Sptan1
Sptbn1
Tnnc1
Tnnt2
Myogenic
c
Myo IIb
regulation of muscle hypertrophy
muscle structure development
muscle cell differentiation
regulation of system process
response to insulin
PI3K-Akt signaling pathway
Insulin secretion
AMPK signaling pathway
Insulin signaling pathway
mTOR signaling pathway
extracellular matrix organization
external encapsulating structure organization
extracellular structure organization
d
Myo IIb → Myo bulk/ Stroma/ Myogenic/ Immune
Myonuclei
Myogenic
Stroma
Immune
Insulin-like growth factor 1
Integrin alpha-8
Bromodomain-containing protein 1
Casp. recruitment domain-containing protein 9
Ciliated left-right organizer metallopeptidase
Microtubule cross-linking factor 2
e
Myonuclei (Myo_I, IIa, IIb, IIxb, New, NMJ)
Myo IIb
Myocytes
Activated MuSC
Trim63
Traf6
Prkn
Usp19
Fbxo30
Fbxo32
Fbxo40
Zfp36l1
Nedd4
Psma1
Eif4ebp1
Gadd45a
Ctsl
Smad3
Tgfbr1
Tgfb3
Mstn
Foxo3
Ip6k1
Park7
Igf1
Irs1
Pik3ca
Akt1
Akt2
Igf1r
Ubiquitin ligases
FoxO/ TGFbeta
IGF pathway
f
Endothelial Cells I
tube morphogenesis
Extracellular matrix organization
Collagen degradation
Degradation of the extracellular matrix
vasculature development
blood vessel morphogenesis
blood vessel development
angiogenesis
PI3K-Akt signaling pathway
Focal adhesion PI3K Akt mTOR signaling pathway
g
Endothelial Cells I
Igf1
Met
Gab1
Calcrl
Mmp14
Col18a1
Col4a1
Col4a2
Nrp2
Plxnd1
Clec14a
Thsd7a
Vav2
Arhgap22
Prcp
Pro-angiogenesis
h
Geriatric plantaris
Sham vehicle
Overload vehicle
Overload PGDHi
CD31/DAPI
i
CD31 area (%)
Sham vehicle
Overload vehicle
Sham veh
Overload vehicle
Overload PGDHi
Young
Geriatric

**Figure 6. PGDHi enhances IGF signaling and vascular growth in overloaded aged muscle.**
(a) Bar graph showing the number of differentially expressed genes (DEGs) over sham in selected cell populations (MuSC, activated MuSC, myocytes, Myo_New, Myo_IIb, Myo_IIa) from overloaded or PGDHi-treated overloaded muscles.
(b) Analysis of the Myo_New population. UMAP visualization of Myo_New nuclei from young vehicle, geriatric vehicle, and geriatric PGDHi groups (left). Gene ontology (GO) terms enriched among DEGs in Myo_New nuclei of overloaded muscles (right heatmap). Dot plot of representative DEGs in Myo_New nuclei across groups (bottom), with dot size indicating fraction of nuclei expressing each gene and color indicating mean expression.
(c) UMAP visualization of Myo_IIb nuclei from young vehicle, geriatric vehicle, and geriatric PGDHi groups (left), with enriched gene ontology (GO) terms shown as a heatmap (right).
(d) Venn diagram of predicted ligand–receptor interactions from Myo_IIb to bulk myonuclei, stromal, myogenic, and immune compartments, highlighting IGF1 as a shared interactor.
(e) Dot plots showing expression of genes related to ubiquitin ligases, FoxO/TGFβ signaling, IGF pathway, and representative targets in myocytes and activated MuSCs across conditions. Dot size indicates the fraction of cells expressing the gene, and color intensity represents mean expression.
(f) UMAP visualization of endothelial cell I populations (left) with enriched GO terms shown as a heatmap (right).
(g) Dot plots of pro-angiogenic DEGs in endothelial cells across conditions.
(h) Representative immunofluorescence images of CD31+ endothelial cells (green) in plantaris muscle sections from geriatric mice under sham, overloaded vehicle, and overloaded PGDHi conditions. Nuclei counterstained with DAPI (blue). Scale bar, 100 μm.
(i) Quantification of CD31+ area as a percentage of total tissue area (n = 8-12 per group). Each dot represents one animal. Data represent mean ± SEM. *$P < 0.05$, **$P < 0.01$, ***$P < 0.001$, **** $P < 0.0001$ by unpaired t-test or one-way ANOVA with post hoc test, as appropriate.

### Figure S5

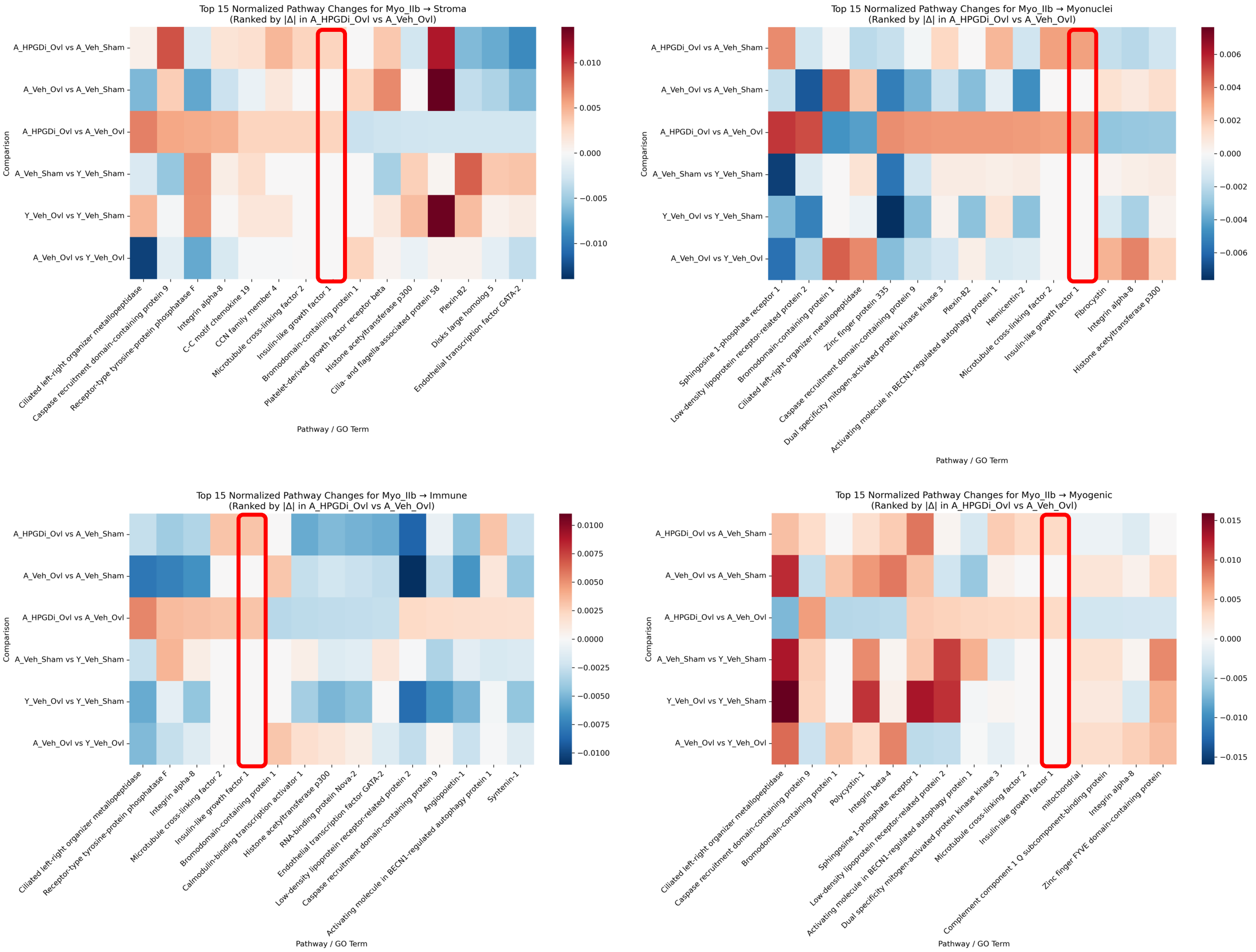


**Figure S5. Analysis of ligand–receptor interactions from Myo_IIb nuclei reveals IGF pathway enrichment.** Heatmaps showing the top 15 normalized pathway changes for Myo_IIb-derived interactions with stromal cells (top left), myonuclei (top right), immune cells (bottom left), and myogenic populations (bottom right). Rows represent pairwise comparisons across groups (young sham, young overload, geriatric sham, geriatric overload, and geriatric overload with PGDHi). Columns indicate enriched GO terms. Color scale reflects normalized pathway activity changes, with red indicating enrichment and blue indicating downregulation.

**Figure 7**

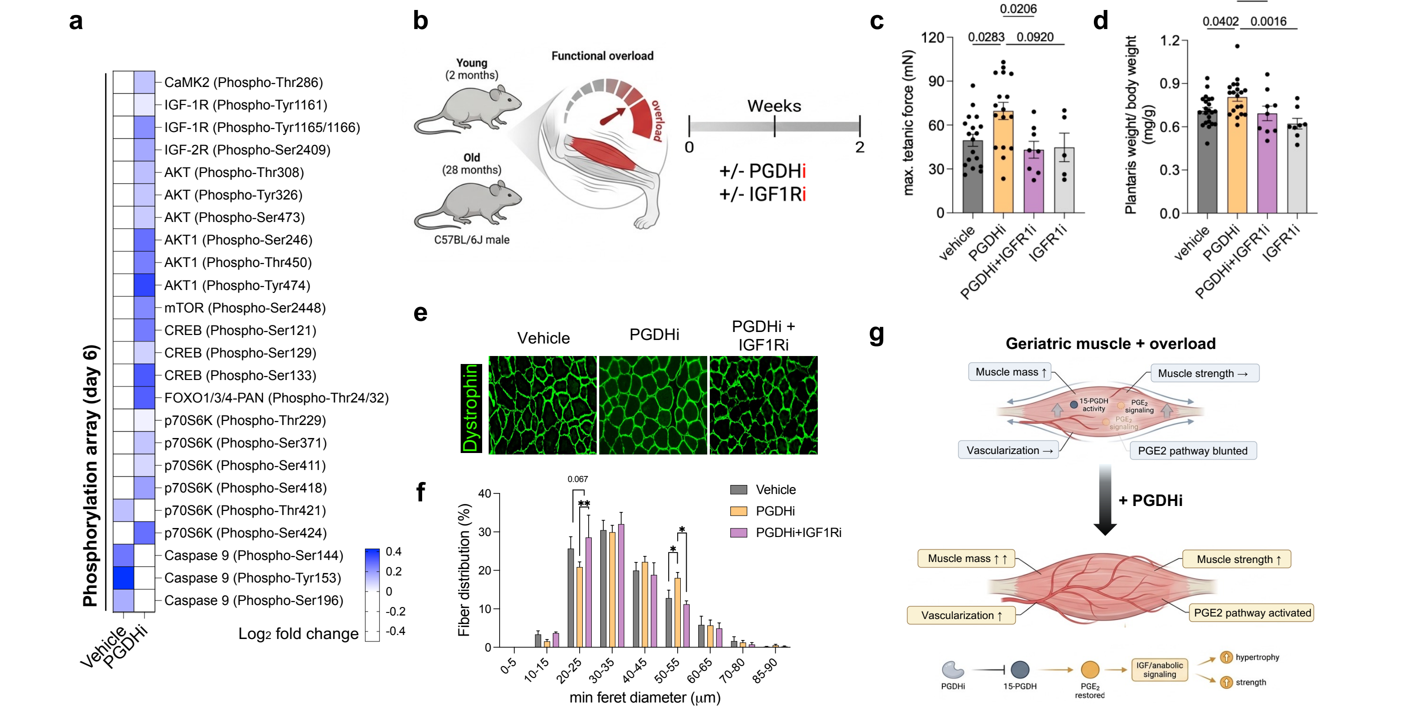


**Figure 7. IGF1 signaling is required for the hypertrophic effects of PGDHi in aged muscle.**
(a) Phosphorylation array of IGF1R-related signaling proteins from overloaded plantaris muscles of geriatric mice treated with vehicle or PGDHi for 6 days. Heatmap shows $\log_2$ fold change relative to vehicle.
(b) Experimental schematic of young and aged mice subjected to overload and treated with PGDHi, with or without IGF1 receptor inhibitor (IGF1Ri), for two weeks.
(c) Absolute plantarflexor force in geriatric mice across treatment groups (n = 5-18 per group).
(d) Plantaris muscle weight in geriatric mice across treatment groups (n = 5-18 per group).
(e) Representative immunofluorescence images of dystrophin-stained muscle cross-sections from geriatric mice treated with vehicle, PGDHi, or PGDHi + IGF1Ri. Scale bar, 100 µm.
(f) Fiber size distribution of plantaris myofibers in geriatric mice treated as in (e).
(g) Schematic summary illustrating impaired overload response in aged muscle (top), and how PGDHi restores hypertrophic adaptations through enhanced PGE2 bioavailability and IGF1 induction in type IIb fibers (bottom).
Data represent mean ± SEM. *$P < 0.05$, **$P < 0.01$, ***$P < 0.001$, **** $P < 0.0001$ by unpaired t-test or one-way ANOVA with post hoc test, as appropriate.

| | | |
|---|---|---|
| Igf1 | Forward | AGCAGCCTTCCAACTCAATTAT |
| | Reverse | GAAGACGACATGATGTGTATCTTTATC |
| Mymk | Forward | ATCGCTACCAAGAGGCGTT |
| | Reverse | CACAGCACAGACAAACCAGC |
| Mymx | Forward | CAGGAGGGCAAGAAGTTCAG |
| | Reverse | ATGTCTTGGGAGCTCAGTCG |
| MyoG | Forward | GAGACATCCCCCTATTTCTACCA |
| | Reverse | GCTCAGTCCGCTCATAGCC |
| Myf5 | Forward | AAGGCTCCTGTATCCCCTCAC |
| | Reverse | TGACCTTCTTCAGGCGTCTAC |
| Psx7 | Forward | TCTCCAAGATTCTGTGCCGAT |
| | Reverse | CGGGGTTCTCTCTCTTATACTCC |
| EP2 (Ptger2) | Forward | TCCCTAAAGGAAAAGTGGGAC |
| | Reverse | GAGCGCATTAACCTCAGGACC |
| EP4 (Ptger4) | Forward | ACCATTCCTAGATCGAACCGT |
| | Reverse | CACACCCCGAAGATGAACAT |
| COX-2 (Ptgs2) | Forward | TGAGCAACTATTCCAAACCAGC |
| | Reverse | GCACGTAGTCTTCGATCACTATC |
| 16s | Forward | CCGCAAGGGAAAGATGAAAGAC |
| | Reverse | TCGTTTGGTTTCGGGGTTTC |
| 15-PGDH (Hpgd) | Forward | GGAAGAGCCGAAATTATTCGCT |
| | Reverse | ACCACTGCATCAGCTTGACAT |

**Table S1.** List of primers used for the real-time qPCR experiments.
